# Chronic contractile activity induced skeletal muscle-derived extracellular vesicles increase mitochondrial biogenesis in recipient myocytes via transmembrane or peripheral membrane proteins

**DOI:** 10.1101/2024.08.01.605916

**Authors:** Patience O. Obi, Tamiris F. G. Souza, Samira Seif, Benjamin Bydak, Nicholas Klassen, Adrian R. West, Joseph W. Gordon, Ayesha Saleem

**Affiliations:** Diabetes Research Envisioned and Accomplished in Manitoba (DREAM) Research Theme, Winnipeg, MB, Canada; Biology of Breathing Research Theme, Winnipeg, MB, Canada; Applied Health Sciences, University of Manitoba, Winnipeg, MB, Canada; Department of Physiology and Pathophysiology, Rady Faculty of Health Sciences, University of Manitoba, Winnipeg, MB. Canada; College of Nursing, Rady Faculty of Health Sciences, University of Manitoba, Winnipeg, MB, Canada; Children’s Hospital Research Institute of Manitoba (CHRIM), Winnipeg, MB, Canada; Faculty of Kinesiology and Recreation Management, University of Manitoba, Winnipeg, MB, Canada

**Keywords:** extracellular vesicles, skeletal muscle cells, myoblasts, myotubes, differential ultracentrifugation, chronic contractile activity, mitochondrial biogenesis

## Abstract

The effect of chronic contractile activity (CCA) on the biophysical properties and functional activity of skeletal muscle extracellular vesicles (Skm-EVs) is poorly understood due to challenges in distinguishing Skm-EVs originating from exercising muscle *in vivo*. To address this, myoblasts were differentiated into myotubes, and electrically paced (3 h/day, 4 days @ 14 V). CCA evoked an increase in mitochondrial biogenesis in stimulated *vs*. non-stimulated (CON) myotubes as expected. EVs were isolated from conditioned media from control and stimulated myotubes using differential ultracentrifugation and characterized biophysically using tunable resistive pulse sensing (TRPS, Exoid), TEM and western blotting. TEM images confirmed isolated round-shaped vesicles of about 30 - 150 nm with an intact lipid bilayer. The mean size of EVs ranged from 98 -138 nm, and was not altered by CCA. Zeta potential and total EV protein yield remained unchanged between groups, and total EV secretion increased after 4 days of CCA. Concomitant analysis of EVs after each day of CCA also demonstrated a progressive increase in CCA-EV concentration, while size and zeta potential remained unaltered, and EV protein yield increased in both CON-EVs and CCA groups. CCA-EVs were enriched with small-EVs *vs*. CON-EVs, concomitant with higher expression of small-EV markers CD81, Tsg101 and HSP70. In whole cell lysates, CD63 and ApoA1 were reduced with CCA in myotubes, whereas CD81, Tsg101, Flotillin-1 and HSP70 levels remained unchanged. To evaluate the functional effect of EVs secreted post-CCA, we treated C2C12 myoblasts with all EVs isolated from CON or CCA myotubes after each day of stimulation, and measured cell count, cell viability, protein yield and mitochondrial biogenesis in recipient cells. There was no effect on cell count, viability and protein yield. Myoblasts treated with CCA-EVs exhibited increased mitochondrial biogenesis as indicated by enhanced MitoTracker Red staining, cytochrome *c* oxidase activity, and protein expression of electron transport chain subunit, CIV-MTCO1. Further, CCA-EV treatment enhanced maximal oxygen consumption rates (OCR), and ATP production in treated myoblasts. This increase in maximal OCR was abrogated when CCA-EVs pre-treated with proteinase K were co-cultured with myoblasts, indicating the pro-metabolic effect was likely mediated by transmembrane or peripheral membrane proteins in CCA-EVs. Our data highlight the novel effect of Skm-EVs isolated post-CCA in mediating pro-metabolic effects in recipient cells and thereby transmitting the effects associated with traditional exercise. Further investigation to interrogate the underlying mechanisms involved in downstream cellular metabolic adaptations is warranted.

## Introduction

Extracellular vesicles (EVs) play a central role in cell-cell communication. EVs are found in all biological fluids including blood, urine, cerebrospinal fluid, saliva, breast milk, amniotic fluid, among others [1]–[3]. EVs have been traditionally divided into three subtypes based on their biogenesis, size, and biological properties. They include exosomes (30-150 nm) formed by the invagination of endosomal membranes leading to the formation of intraluminal vesicles enclosed within multivesicular bodies (MVBs) that are subsequently exocytosed through the plasma membrane; microvesicles (100-1000 nm) formed by the outward budding of the plasma membrane, and apoptotic bodies (500-5000 nm) formed from the outward blebbing of an apoptotic cell membrane [4]. The Minimal Information for Studies of EVs (MISEV) guidelines recommend classifying EVs based on i) size, where small EVs (sEVs) are <200 nm and medium/large EVs (m/lEVs) >200 nm, ii) biochemical composition, and iii) cell of origin [3]. EVs contain biological cargo such as DNA, mRNA, miRNA, proteins, lipids, and metabolites and can transfer them from parent to recipient cells [5]. In the last two decades, EVs have been extensively studied as potential biomarkers for many chronic diseases [6]–[10], and as novel therapeutics for drug delivery [11]–[13]. More recently, researchers have been begun to investigate the functional effect of EVs in physiological contexts.

Regular exercise is an important physiological stressor that has a plethora of positive systemic benefits such as improved mitochondrial content and function termed mitochondrial biogenesis, enhanced antioxidant profile, protection against cardiovascular diseases, and improved muscle strength, among many other beneficial adaptations [14]–[16]. Exercise affects almost all the tissues in the body but the key organ systems that beneficially adapt include skeletal muscle, adipose tissue, brain, bone marrow, heart, gut, pancreas, and liver [17]. Skeletal muscle is the largest organ system in the body, accounting for ∼40% of body mass, and is a critical regulator of whole-body metabolic capacity. Strong evidence suggests that the systemic adaptative effects of exercise are potentiated in part by the release of factors known as myokines during exercise from skeletal muscle [15], [18]. Myokines are usually proteins that can travel to different organs in the body and act in an autocrine, paracrine or endocrine manner. Examples include interleukin (IL)-6, IL-10, IL-15, irisin, meteorin, myostatin, among others [18]. In addition to myokines, EVs can also be released from the skeletal muscle during exercise. There is growing interest in how acute and chronic exercise affects EVs and whether they play a role in mediating the beneficial systemic effects of exercise [19]–[22].

Previous studies have shown that acute/chronic exercise in humans, mice and rats increases the release of EVs in the circulation, and modifies EV cargo components [23]–[34]. However, the majority of this work has been done using EVs isolated from whole blood. Whole blood contains a heterogenous mixture of vesicles derived from platelets, erythrocytes, endothelial cells, leukocytes, [24], [27], and with a comparatively small proportion of EVs from skeletal muscle [21]. Given the challenge in differentiating skeletal muscle-derived EVs (Skm-EVs) in systemic EV preparations, and the fact that skeletal muscle is: 1) the major tissue system involved in muscle contraction during physical activity, 2) can modify paracrine and exocrine tissue/organ metabolism through release of myokines directly or packaged in EVs, and 3) accounts for ∼40% of the body mass, it is critical to distill the impact of chronic exercise on the characteristics and biological activity of Skm-EVs specifically. Some have successfully used alpha-Sarcoglycan (SGCA), a protein marker specific to skeletal muscle to distinguish Skm-EVs in whole blood, and found SGCA+ EVs comprised ∼1-5% of total circulatory EVs [21]. While SGCA+ is specific for Skm-EVs, it does not guarantee that all EVs released from skeletal muscle would express it. Another recent study used a skeletal muscle myofiber-specific dual fluorescent reporter mouse to demonstrate the presence of Skm-EVs *in vivo*, and presented corroborating evidence that Skm-EVs constituted ∼5% of circulating EVs [35]. While both studies present viable systems to investigate Skm-EVs *in vivo*, neither model permits distinguishing Skm-EVs originating from the contracting skeletal muscle fibers *vs*. other skeletal muscle fibers at rest. This, in tandem with the reported low concentration of Skm-EVs in circulation, precludes a thorough characterization of the effects of chronic exercise on Skm-EVs and their functional activity, thereby necessitating the use of alternate *in vitro* model system approaches as exemplified by this current work.

In this study, we used chronic electrical pulse stimulation (IonOptix C-PACE EM) to mimic exercise *in vitro*. While chronic electrical pulse stimulation of differentiated skeletal muscle myotubes (immature muscle fibers) may not recapitulate all facets of endurance exercise training, it is a validated and well-established model to study the role of skeletal muscle contraction in exercise-mediated adaptations such as mitochondrial biogenesis. Indeed chronic contractile activity (CCA) *in vitro* can evoke similar adaptations in mitochondrial biogenesis as weeks of exercise training *in vivo* [36], [37]. Using this well-characterized model, we measured the effect of CCA on the release and biophysical characteristics of Skm-EVs, and sought to determine the biological functional activity of Skm-EVs derived post-CCA. We hypothesized that Skm-EVs isolated post-CCA will potentiate an increase in mitochondrial biogenesis in skeletal muscle cells. To test our hypothesis, we purified EVs from conditioned media from non-contracted control myotubes, and from myotubes post-CCA. Both control and CCA-EVs were co-cultured with myoblasts and the subsequent effect on mitochondrial biogenesis measured in recipient cells.

## Methods

### Cell culture

Murine C2C12 myoblasts (500,000 cells/well) were seeded in a six well plate pre-coated with 0.2% gelatin, and grown in fresh Dulbecco’s Modification of Eagle’s Medium (DMEM; Sigma-Aldrich) supplemented with 10% fetal bovine serum (FBS; Gibco/Thermo Fisher Scientific) and 1% penicillin/streptomycin (P/S) (growth media). Cells were grown at 37 °C in 5% CO_2_ incubator for 24 h. When myoblasts reached approximately 90-95% confluency, the growth media was switched to differentiation media (DMEM supplemented with 5% heat-inactivated horse serum (HI-HS; Gibco/Thermo Fisher Scientific) and 1% P/S) for 5 days to differentiate myoblasts into immature muscle fibers, commonly referred to as myotubes.

### Study design: chronic contractile activity (CCA) and sample collection for EV purification

Chronic contractile activity (CCA) was performed as previously described by Uguccioni and Hood [36] with some modifications. After 5 days of differentiation, myotubes were divided into control (CON) and CCA groups. The CCA plates were stimulated using the C-Pace EM cell culture stimulator, with C-dish and carbon electrodes (IonOptix, Milton, MA, United States) while the CON plates had the C-dish placed on top but without the carbon electrodes. Myotubes were subjected to CCA at 1 Hz (2 ms pulses), 14 V for 3 h/day for 4 consecutive days. After each bout of contractile activity, spent media was discarded and 2 mL fresh differentiation media added to each well. This is important as previous observations by Saleem and Hood (unpublished, PhD dissertation) indicate that changing spent media post-contractile activity improves protein and RNA yield from CCA myotubes. Myotubes were left to recover for 21 h before the next bout of contractile activity. On day 4, immediately after the last bout of contractile activity, media was changed to exosome-depleted differentiation media (DMEM supplemented with 5% exosome-depleted HI-HS and 1% P/S) for both groups and myotubes left to recover for 21 h. Exosome-depleted horse serum was used to remove any confounding effects of EVs found naturally in horse serum. To deplete exosomes from horse serum, we diluted 5 mL of HI-HS to 10% using DMEM, spun at 100,000*xg* for 18 h and filtered with 0.22 µm filter. This exosome-depleted HI-HS was then used to make the exosome-depleted differentiation media using standard formulation as described above. After 4 days of CCA, post-21 h recovery, 12 mL conditioned media from CON and CCA myotubes was collected and used for EV isolation and characterization as described below. Control and CCA myotubes were harvested for MitoTracker Red CMXRos staining, COX activity and western blotting assays to confirm CCA elicited an increase in mitochondrial biogenesis in stimulated myotubes.

A separate set of experiments were done to evaluate the effect of each bout of contractile activity on Skm-EVs. Here, after day 1 of contractile activity, media was changed to exosome-depleted differentiation media and myotubes were left to recover for 21 h. Afterwards, conditioned media was collected and used for EV isolation. This process was repeated after each day for 4 consecutive days. CCA-EVs collected after the first bout of contractile activity were labelled Day 1 EVs, after the second day of CCA as Day 2 EVs and so on till Day 4. CON-EVs were collected from Day 1 – Day 4 in parallel.

### Isolation of EVs by differential centrifugation (dUC)

EV isolation by differential ultracentrifugation (dUC) was performed according to the protocol by Théry et al. [38]. Briefly, conditioned media (CM) from each six well plate (12 mL) was collected. For EV-treatment experiments, 2 mL of CM from each condition was kept aside to be used as a positive control. For EV characterization experiments, all 12 mL were utilized for EV isolation. The collected CM was centrifuged at 300*xg* for 10 min at 4 °C to pellet dead cells (Sorvall™ RC 6 Plus Centrifuge, F13-14 fixed angle rotor), followed by centrifugation at 2000*xg* for 10 min at 4 °C to remove cell debris. The resulting supernatant was then centrifuged at 10,000*xg* for 30 min at 4 °C to remove large vesicles. Using an ultracentrifuge (Sorvall™ MTX 150 Micro-Ultracentrifuge, S58-A fixed angle rotor), the supernatant was then spun at 100,000*xg* for 70 min at 4 °C to obtain the exosome-enriched sEV pellet. The supernatant from this step was collected and stored as EV-depleted conditioned media (EV-dep) for later use as a negative control in EV-treatment experiments. The sEV pellet was resuspended in 1 mL PBS and centrifuged again at 100,000*xg* for 70 min at 4 °C. After centrifugation, the final sEV pellet was resuspended in 50 µL PBS and used for subsequent analysis or functional assays.

### Tunable resistive pulse sensing (TRPS)

Concentration, size distribution and zeta potential of isolated EVs (purified 4 days post-CCA, as well as the vesicles purified after each day of CCA) were determined by the orthogonal tunable resistive pulse sensing (TRPS) technology (Exoid; Izon Science Ltd, Christchurch, New Zealand). EV samples were diluted with filtered measurement electrolyte solution (Izon Science Ltd.) and passed through 0.22 µm filters before using for characterization. Calibration particles (Izon Science Ltd.) with standardized diameters made from carboxylated polystyrene beads were also diluted with measurement electrolyte solution. Data were recorded and analyzed using Izon Control Suite software v3.4 (Izon Science Ltd.). Two to four samples were measured at a time, and a calibration was performed after the first sample run using 200 nm calibration particles. Purified EVs were analyzed for size and concentration using NP150 nanopores (Izon Science Ltd.). All calibrations and sample measurements were run under the same conditions recommended by the manufacturer and a minimum of 500 particles was recorded at three different pressures. For size and zeta potential measurement, EV samples were diluted but not filtered, and calibration run was performed using either 100 nm or 200 nm calibration particles.

### Transmission electron microscopy (TEM)

Freshly isolated EVs (10 µl) were fixed with equal volume of 4% paraformaldehyde and carefully applied onto a carbon grid for 20 min at room temperature. Grids were washed with 100 µL PBS and fixed with 1% glutaraldehyde for 5 min. To remove the excess fixation buffer, the grids were washed 8 times with 100 µL distilled water, after which they were negatively stained with 1% uranyl acetate for 2 min and left to airdry at room temperature. TEM grids were visualized using Phillips CM10 Electron Microscope operated at 60 kV using the fee-for-service at the Histology core facility, Department of Human Anatomy and Cell Science, University of Manitoba.

### EV protein extraction, concentration and yield

EV preparations were lysed using 1:1 Pierce RIPA solution with protease inhibitor tablet (Roche) and EV concentration was determined using Pierce™ MicroBCA protein assay kit (Thermo Fisher Scientific) following the manufacturer’s instructions. Briefly, the working reagent was prepared by mixing 25 parts of Reagent A, 24 parts of Reagent B and 1 part of Reagent C as detailed before [33]. Standards were prepared by serial dilution of 2 mg/mL bovine serum albumin (BSA) ampule into clean vials using ultrapure water. 150 µL of each standard or sample was added to a 96-well microplate in duplicates, followed by the addition of 150 µL of working reagent to each well. Samples were incubated 37 °C for 2 h and absorbance was measured at 562 nm using a microplate spectrophotometer (BioTek, Epoch). EV protein yield was determined by multiplying the protein concentration (µg/µL) with the total volume (µL) of EV lysate.

### EV fluorescent labeling

For confirmation of EV uptake in cells, EVs were labelled with MemGlow™ 488 Fluorogenic Membrane Probe (Cytoskeleton) [39]. MemGlow was dissolved as 20 µM stock solutions in dimethyl sulfoxide (DMSO) and diluted in PBS to a final concentration of 200 nM. EVs were incubated with 200 nM MemGlow labelling solution for 1 h at room temperature keeping it protected from light. To remove unbound dye from the solution, EVs were either diluted with filtered PBS and centrifuged at 100,000*xg* for 70 min at 4 °C or loaded onto a 35 nm qEVsingle SEC column (Izon Science Ltd.), after which the final labelled EVs were collected. Simultaneously, dye control (200 nM MemGlow in an equivalent volume of PBS) and unstained EV control (EVs without MemGlow) were processed in the same manner. Flow cytometry (Attune NxT Acoustic Focusing Cytometer, Thermo Fisher scientific, Massachusetts, United States) was performed after labelling to determine the presence and percentage of fluorescently labelled EVs. Labelled EVs were diluted 1:3 in 0.22 µM filtered PBS, and MemGlow 488 signal was detected using the Alexa Fluor 488 channel on the flow cytometer. To perform EV cell uptake experiments, 50 µL of EVs (from 10 mL of conditioned media) were labelled with either 200 nM MemGlow 488 or 2 µM PKH67 dye (Sigma-Aldrich) as detailed before [40], and co-cultured with 10,000 myoblasts/well in an 8-well chamber slide for 24 h. Subsequently, cells were fixed with 3.7% paraformaldehyde (PFA), stained with 66 µM of Rhodamine phalloidin and 2-(4-amidinophenyl)-1H-indole-6-carboxamidine (DAPI), and then used for confocal imaging (Zeiss confocal microscope, St. Louis, USA).

### CON-EV and CCA-EV co-culture with C2C12 myoblasts

We evaluated the functional effects of control and CCA-EVs on cell viability and mitochondrial biogenesis in recipient murine myoblasts. To do so, 90,000 murine myoblasts/well were seeded in 6-well plates in 2 mL of growth media and allowed to adhere for 4 h. After the myoblasts had adhered, cells were washed with PBS and 2 mL of exosome-depleted differentiation media was added to cells. There were three main treatment groups in each experiment: 1) 2 mL CM, experimental control for control and CCA conditions; 2) 2 mL EV-dep CM, negative control for CON-EVs and CCA-EVs; and 3) CON-EVs and CCA-EVs treatments. All EVs isolated each day from 10 mL of conditioned media were used to treat the cells, to recapitulate the physiological relevance of secreted EVs isolated after each bout of contractile activity. Treatment was done using media collected 21 h after each day of contractile activity, meaning on the first day of treatment, Day 1 EVs, CM or EV-dep were added to the cells and left to incubate for 24 h, and this was done continuously for 4 days after each bout of CCA. After the last day of treatment, media was discarded and treated myoblasts were harvested for protein extraction, yield, cell count and viability analyses, and assessment of mitochondrial biogenesis using MitoTracker Red CMXRos staining, COX activity, western blotting and oxygen consumption rate analysis as described below.

### Cell enzyme extraction

Myotube or EV-treated myoblasts whole cell extracts were harvested as previously described by Uguccioni and Hood [36] with some modifications. Cells were harvested using a cell scraper (Corning®), resuspended in 50 µL of enzyme extraction buffer (100 mM Na-K-phosphate, 2 mM EDTA, pH 7.2), sonicated 3 x 3 secs on ice, subjected to repeated freeze-thaw cycles, vortexed vigorously for 5 secs and centrifuged at 14,000 rpm. The supernatant was collected and used to measure protein concentration and yield, COX enzyme activity and western blotting.

### Cell protein concentration and yield

Protein concentration of myotube lysates or EV-treated myoblast whole cell lysates was determined using Pierce™ BCA protein assay kit (Thermo Fisher Scientific) following the manufacturer’s instructions as described before [41]. Briefly, a serial dilution of 2 mg/mL bovine serum albumin (BSA) in sterile water was prepared and used as standard. 25 µL of each standard or sample was added in duplicates into a 96-well microplate, followed by adding 200 µL of the working reagent to each well. Samples were incubated at 37 °C for 30 min and the absorbance was measured at 562 nm using a microplate spectrophotometer (BioTek, Epoch). Protein yield was determined by multiplying the protein concentration (µg/µL) with the total volume (µL) of cell lysates.

### Cell count and viability via trypan blue exclusion assay

Cell count and viability was measured with trypan blue exclusion as described previously with few modifications [42]. Treated myoblasts were washed twice with 1 mL PBS, trypsinized with 500 µL of trypsin, and incubated for 3 min at 37 °C. 1 mL of growth media was added, and cells were centrifuged at 1000*xg* for 5 min to pellet the cells. Pelleted cells were stained with a 1:25 dilution of Trypan blue (Sigma-Aldrich Co., cat #T8154, Saint Louis, MO, United States) then counted with a hemocytometer (Hausser Scientific Bright-Line Hemacytometer, Sigma-Aldrich Co., Saint Louis, MO, USA). Total number of cells were counted and expressed per mL of growth media. Cell viability was obtained by dividing the number of live cells by total number of counted cells.

### Cell viability via MTT assay

Cell viability was also measured by Thiazolyl Blue Tetrazolium Bromide (MTT; Sigma, St. Louis, MO) assay. C2C12 myoblasts were seeded at 5000 cells/well in 96-well plates in triplicates, and treated with CON-EVs or CCA-EVs for 4 days after each bout of contractile activity as described above. After treatment, 20 µL of MTT Reagent 5 mg/mL was added and left for 3 h. The yellow tetrazolium MTT was reduced to formazan crystals by intracellular NAD(P)H-oxidoreductases. 150 µL of DMSO was added and mixed gently to solubilize the formazan crystals. The assay was quantified by spectrophotometry at the absorbance of 540 nm (Agilent BioTek Epoch, Santa Clara, CA, United States).

### MitoTracker Red staining

MitoTracker Red stains mitochondria in live cells and is used as a proxy for mitochondrial mass [37], [43]. Myotubes collected after last bout of contractile activity as well as EV-treated myoblasts, were incubated with MitoTracker Red CMXRos (Cell Signaling Technology) and Hoechst 33258 (Sigma-Aldrich) at final concentrations of 50 nM and 3 µg/mL respectively under normal culture conditions for 30 min. Following incubation, cells were washed with PBS, and medium was switched to differentiation or growth media. Cells were visualized using the Olympus IX70 inverted microscope (Toronto, ON, Canada) with NIS Elements AR 3.0 software (Nikon Instruments Inc., Melville, NY, USA). Two to three representative images per well were taken, and the fluorescence intensity was quantified using ImageJ software (NIH, Bethesda, MD, USA), where the free hand tool was used to draw the region of interest (ROI) from at least three myotubes or five myoblasts for each representative image, and mean intensity for each was averaged to get the total fluorescence intensity.

### COX activity

We measured mitochondrial cytochrome *c* oxidase (COX) activity, a gold standard marker of mitochondrial biogenesis, in myotube or treated myoblast enzyme extracts as previously described [44]. Briefly, enzyme extracts were added to a test solution containing fully reduced cytochrome *c* (Sigma-Aldrich). COX activity was determined as the maximal rate of oxidation of fully reduced cytochrome *c* measured by the change in absorbance over time at 550 nm using a cell imaging multi-mode reader (Agilent BioTek Cytation 5, Santa Clara, CA, United States) at 30 °C. Protein concentration was determined using the Pierce™ BCA protein assay kit as described above, and COX activity was normalized by total protein content.

### Western blotting

5 µg of total protein from EVs or whole cell extracts were resolved on a 12% SDS-PAGE gel and subsequently transferred to nitrocellulose membranes. Membranes were then blocked for 1 h with 5% skim milk in 1× Tris-buffered saline-Tween 20 solution (TBST) at room temperature, followed by incubation with primary antibodies in 1% skim milk overnight at 4 °C. The following primary antibodies were used: rabbit polyclonal anti-CD63 (SAB4301607, Sigma-Aldrich, 1:200), mouse monoclonal anti-CD81 (sc-166029, Santa Cruz Biotechnology, 1:200), rabbit polyclonal anti-Tsg101 (T5701, Sigma-Aldrich, 1:1000), rabbit polyclonal anti-Flotillin-1 (F1180, Sigma-Aldrich, 1:200), mouse monoclonal anti-heat shock protein 70 (HSP70) (H5147, Sigma-Aldrich, 1:500), rabbit monoclonal anti-Alix (MCA2493, Bio-Rad Laboratories, 1:200), mouse monoclonal anti-Apolipoprotein A1 (ApoA1) (0650-0050, Bio-Rad Laboratories, 1:200), rabbit polyclonal anti-cytochrome *c* (AHP2302, Bio-Rad Laboratories, 1:500), and mouse monoclonal anti-β-actin (A5441-.2ML, Sigma-Aldrich, 1:5000), mouse oxidative phosphorylation [OXPHOS] cocktail (45-8099, Invitrogen, 1:500) and rabbit monoclonal anti-TFAM (8076, Cell Signaling, 1:200). Subsequently, membranes were washed three times for 5 min with TBST, followed by incubation with anti-mouse (A16017, Thermo Fisher) or anti-rabbit (A16035, Thermo Fisher) IgG horseradish peroxidase secondary antibody (1:1,000-10,000) in 1% skim milk for 1 h at room temperature, after which the membrane was washed again 3 times x 5 min each with TBST. Membranes were developed using enhanced chemiluminescence detection reagent (Bio-Rad Laboratories), and the films were scanned using the ChemiDoc^TM^ MP Imaging System (Bio-Rad Laboratories, Hercules, CA, United States). The intensity of bands was quantified using the Image Lab Software (Bio-Rad Laboratories) and corrected for loading using Coomassie blue or Ponceau S staining.

### Oxygen consumption rate (OCR) using Seahorse XFe24 Analyzer

Oxygen consumption rate (OCR) in EV-treated myoblasts was measured with a Seahorse Flux Analyzer XFe24 (Agilent Technologies, Santa Clara, CA, United States) using the Cell Mito Stress Test Kit (Agilent Technologies) according to the manufacturer’s recommendation and as described before [45], with some modifications. Briefly, 24 h after the last day of EV treatment, myoblasts were trypsinized and seeded (10,000 cells/well) in triplicates per condition in a Seahorse XFe24 plate and cultured overnight. The next day, cells were washed with Seahorse XF DMEM Medium (pH = 7.4) supplemented with XF glucose (10 mM), XF pyruvate (1 mM), and XF glutamine (2 mM), and were incubated in a CO_2_-free incubator at 37 °C for 1 h. The same Seahorse XF DMEM media was used to prepare the injection compounds: i) oligomycin (complex V inhibitor, 1.5 µM), ii) carbonyl cyanide-p-trifluoromethoxyphenylhydrazone (FCCP, uncoupling agent, 2 µM), and iii) rotenone/antimycin A (complex I and complex III inhibitors, respectively, 0.5 µM each). Three measurements of OCR and extracellular acidification rate (ECAR) were recorded before and after the sequential injection of the compounds. Data were normalized for protein yield. The basal OCR was derived by subtracting the baseline rate from rotenone/antimycin A rate; maximal OCR by subtracting FCCP rate from rotenone/antimycin A rate; ATP production by subtracting oligomycin rate from baseline rate; proton leak by subtracting oligomycin rate from rotenone/antimycin A rate; spare capacity by subtracting maximal respiration from basal respiration; and non-mitochondrial respiration is the rotenone/antimycin A rate.

### EV protein digestion with Proteinase K and Triton X-100

To determine whether the effect of CCA-EVs on mitochondrial biogenesis was mediated by lumenal *vs*. membrane-bound or extramembranous protein cargo, isolated EVs were treated with or without 0.1% Triton X-100 at room temperature, and incubated with 10 µg/mL proteinase K (ProK; Sigma Aldrich) for 1 h at 37 °C. ProK activity was inhibited by adding 5 mM phenylmethylsulfonyl fluoride (PMSF; Sigma Aldrich) for 10 min at room temperature. The samples were further processed by addition of 6 mL PBS followed by ultracentrifugation at 100,000*xg* for 70 min at 4 °C to eliminate Triton before co-culture with cells. After centrifugation, the final pellet (not visible) was resuspended in 50 µL PBS, and either characterized using TRPS and western blot or co-cultured with myoblasts as described previously. For co-culture experiments, only CCA-EVs were pretreated with or without Triton X-100 and ProK before co-culture with myoblasts. Myoblasts were treated for 4 days as previously described, and after the last day of treatment, cells were analyzed for OCR using Seahorse assay as described above.

### Statistical analyses

Data were analyzed using unpaired Student’s t-test, one-way ANOVA, and/or two-way ANOVA. Multiple comparisons in the one-way ANOVA were corrected using Holm-Šídák post hoc test, or if the data did not pass normality, with Dunn’s post hoc test. Normality was measured using a combination of tests: D’Agostino-Pearson, Anderson-Darling, Shapiro-Wilk and Kolmogorov-Smirnov when possible. Multiple comparisons in the two-way ANOVA were corrected using Bonferroni’s post hoc test. Individual data points are plotted, with mean ± standard error of mean (SEM) shown as applicable. All graphs were created using GraphPad-Prism software (version 10.1.2, GraphPad, San Diego, CA, USA). Significance was set at p ≤ 0.05. Exact p values for significant or close to statistically significant results is shown. A sample size (n) = 3-9 was conducted for all experiments.

## Results

### CCA evokes mitochondrial biogenesis in C2C12 myotubes

As illustrated in **Fig. 1A**, we first sought to validate the model of chronic contractile activity (CCA) used, by examining key markers of mitochondrial biogenesis in contracted myotubes. Control and CCA myotubes were stained with 50 nM MitoTracker CMXRos for 30 min at 37 °C after conditioned media was collected for EV isolation. Representative fluorescent images at 40X and 10X are shown for CON and CCA myotubes (**Fig. 1B**). Quantification of MitoTracker Red staining showed a 1.1-fold increase in CCA *vs*. CON myotubes (p=0.0445, n=8, **Fig. 1B**). In tandem, we observed a 1.5-fold increase in COX activity with CCA (p=0.0192, n=8, **Fig. 1C**), indicative of an increase in mitochondrial biogenesis as previously detailed [36], [46]. We also evaluated the expression of cytochrome *c* (Cyt C; a mitochondrial protein important in electron transport chain) in CCA *vs*. CON myotubes using western blotting. CCA myotubes showed a 2.2-fold increase in expression of Cyt C relative to CON myotubes (p=0.0120, n=7, **Fig. 1D**). Lastly, we measured protein yield, which was unchanged between groups (**Fig. 1E**).

**Figure 1.**
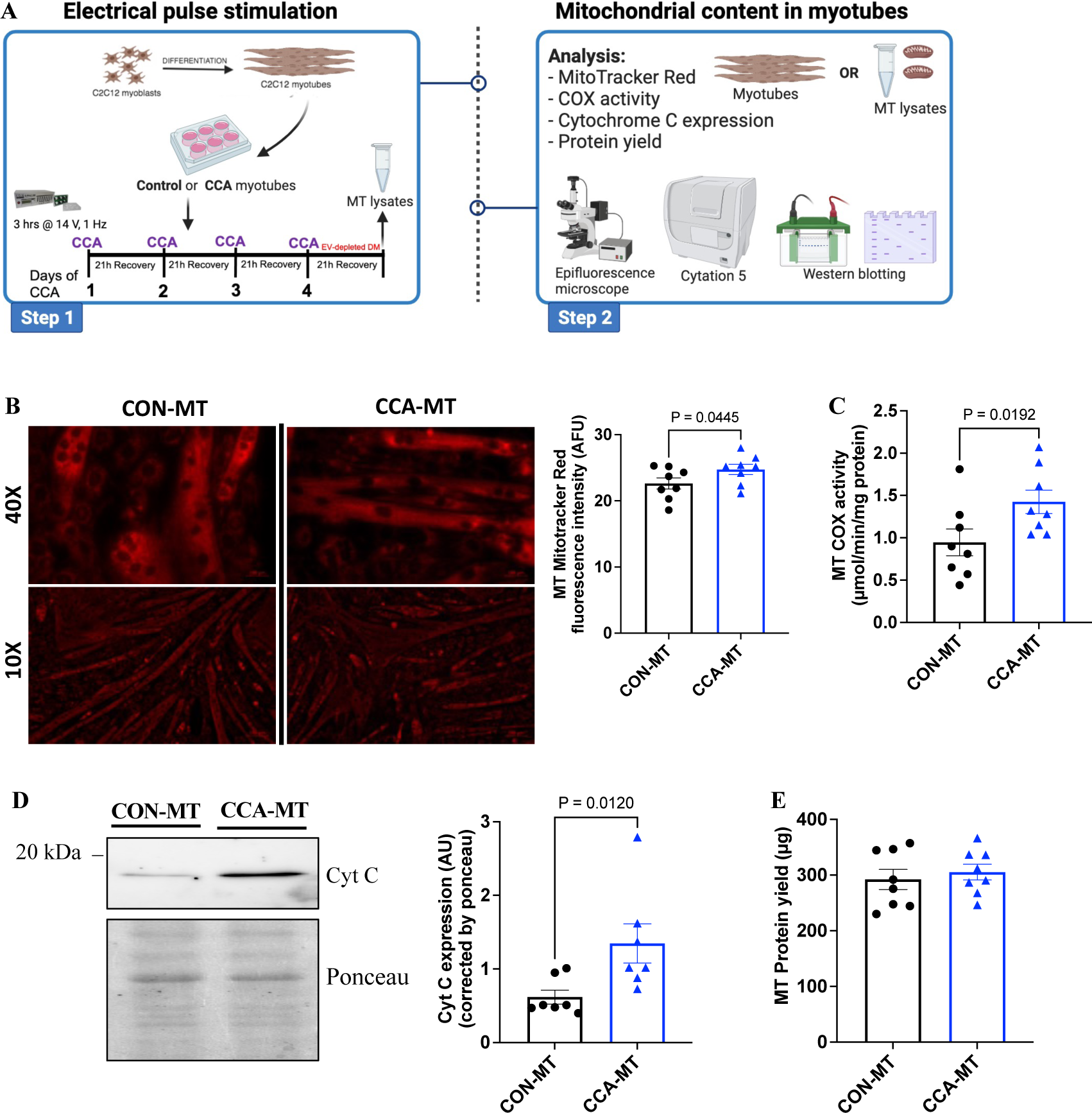
CCA evokes mitochondrial biogenesis in C2C12 murine myoblasts. **(A)** C2C12 myoblasts were fully differentiated into myotubes (MTs). MTs were divided into control (CON) and chronic contractile activity (CCA) plates. CCA-MTs were electrically paced for 3h/day x 4 days at 14 V to mimic chronic endurance exercise *in vitro* using IonOptix ECM. Media was changed after each day of contractile activity and cells were left to recover for 21 h after each bout of contractile activity. After the last day of CCA, media was switched to differentiation media with exosome-depleted horse serum in both CON and CCA-MTs and cells were allowed to recover for 21 h. Conditioned media from control or stimulated myotubes was collected and MT lysates were harvested for mitochondrial biogenesis measurement. **(B)** Representative fluorescent images of CON-MT and CCA-MT stained with MitoTracker Red at 40X and 10X magnification with the graphical quantification on the right, scale bar=100 µm. **(C)** Cytochrome *c* oxidase (COX) activity, **(D)** cytochrome *c* (Cyt C) expression and **(E)** protein yield in CCA-MTs *vs*. CON-MTs. Data were analyzed using an unpaired Student’s t-test and expressed as scatter plots with mean (n=7-8). Exact p values for significant results (p<0.05) are shown. Figure 1A created with BioRender.com.

### CCA increases sEV secretion

As graphically illustrated in **Fig. 2A**, we subsequently evaluated the effects of CCA on Skm-EVs by isolating and characterizing the vesicles biophysically. The protocol for depletion of exosomes/sEVs from horse serum (**Fig. S1A**) was effective as demonstrated by a lack of EVs in the depleted samples (**Fig. S1B**). EVs were isolated from conditioned media of CON and CCA myotubes after 4 days of CCA. Morphological analysis using TEM revealed that myotubes released spherical EVs of about 100 nm with intact double lipid layer, indicating presence of sEVs (**Fig. 2B**). Using TRPS, the orthogonal single-particle measurement technique for nanovesicles, we characterized EVs by size, concentration, and zeta potential. The average size of EVs was similar between CON-EVs (116.3 nm) and CCA-EVs (119.4 nm) (**Fig. 2C**). The average concentration of CCA-EVs (2.26E10 particles/mL) was 1.7-fold higher than CON-EVs (1.26E10 particles/mL) (p=0.0149, n=6, **Fig. 2D**). Size distribution histogram also reflected elevated CCA-EV concentration, with a specific enrichment of sEVs in CCA-EVs *vs*. CON-EVs (**Fig. 2E**). Zeta potential and EV protein yield were not different between the two groups (**Fig. 2F** and **Fig. 2G**). To complement the TRPS findings, we used DLS to measure EVs and found that CCA-EVs are enriched with sEVs (< 200 nm) compared to CON-EVs (**Fig. S2**). Overall, our results indicate that 4 days of CCA increased the secretion of sEVs from myotubes.

**Figure 2.**
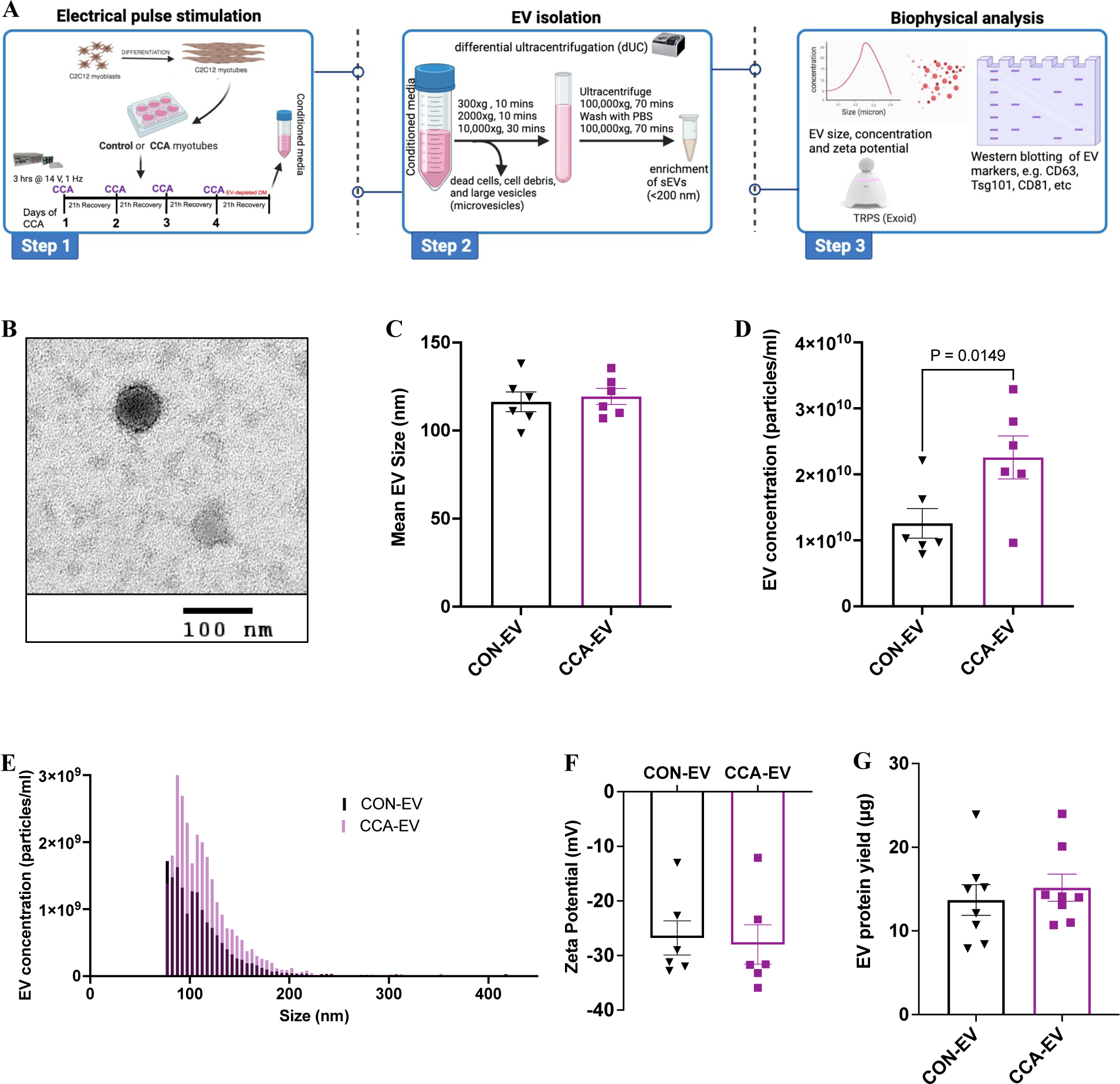
CCA increases small EV secretion from myotubes. **(A)** CCA was done as previously described. Conditioned media from control or stimulated myotubes was collected. EVs were isolated via differential ultracentrifugation (dUC) and characterized by size, zeta potential and concentration using tunable resistive pulse sensing (TRPS). Protein yield and markers of EV subtypes were also measured. (**B)** Representative TEM images of EVs released from myotubes shows the presence of sEVs around 100 nm in diameter: scale bar=100 nm. **(C)** Average EV size, **(D)** EV concentration, **(E)** EV size/concentration histogram, **(F)** zeta potential and **(G)** EV protein yield in CCA-EVs *vs*. CON-EVs. Data were analyzed using an unpaired Student’s t-test in panels C, D, F and G (n=6-8). Exact p values for significant results (p<0.05) are shown. Figure 2A created with BioRender.com.

### CCA alters proteins associated with EV subtypes

As per MISEV guidelines [47], we evaluated expression of protein markers of EV origin, size (sEVs or m/lEVs) and purity of isolation (co-isolated non-EV proteins). We measured proteins normally enriched in exosomes or sEVs (CD81, CD63, Tsg101, HSP70 and Flotillin-1, Alix), proteins found in m/lEVs (Cyt C and β-actin), and a non-EV marker (Apo-A1). Interestingly, CCA-EVs showed increased expression of CD81 (by 1.4-fold, p=0.0465, n=9), Tsg101 (by 1.2-fold, p=0.0497, n=9), and HSP70 (by 1.9-fold, p=0.0227, n=9) (**Fig. 3A** and **3B**) compared to CON-EVs. We found no difference in the expression of CD63, Flotillin-1, Alix and ApoA1 between the two groups, and could not detect any expression of Cyt C and β-actin in the EV preparations from either group (**Fig. 3A** and **3B**). Overall Skm-EVs generally express markers of sEVs, with an enrichment of CD81, Tsg101, and HSP70 in CCA-EVs, indicating an increase in sEVs with chronic contractile activity corroborating the size and concentration results mentioned earlier. We also measured the expression of CD81, CD63, Tsg101, HSP70, Flotillin-1, and ApoA1 in CON and CCA myotube whole cell lysates to deduce if CCA affected proteins involved in EV biogenesis and secretion. Surprisingly, CCA myotubes showed decreased expression of CD63 (by 1.5-fold, p=0.0328, n=7) and ApoA1 (by 3.1-fold, p=0.0261, n=7) compared to CON myotubes (**Fig. 4A** and **4B**). No differences in the expression of CD81, Tsg101, HSP70 and Flotillin-1 between CON and CCA myotubes were observed (**Fig. 4A** and **4B**).

**Figure 3.**
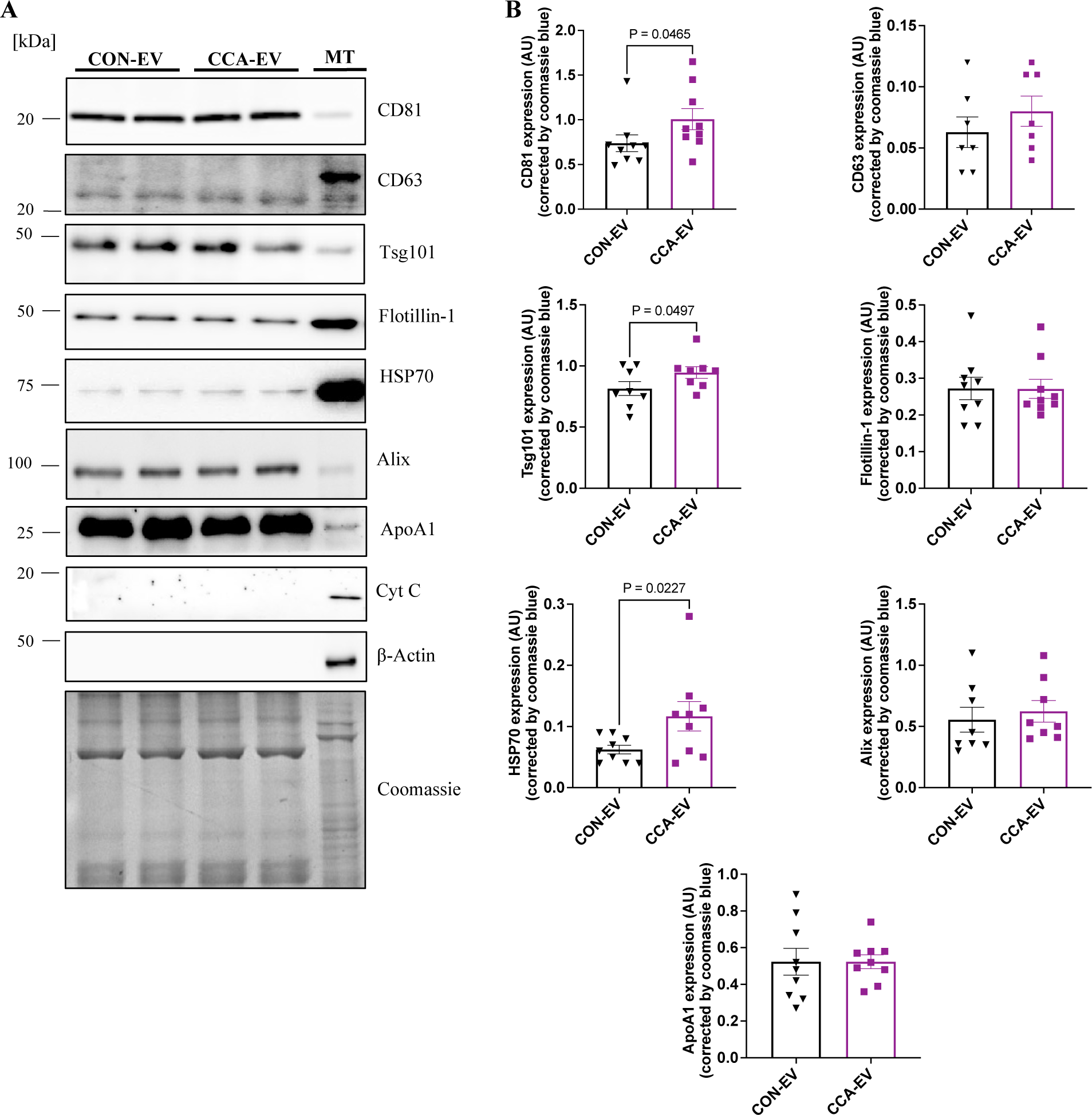
CCA alters expression of proteins related to EV subtypes. **(A)** Equal amounts (5 µg) of proteins from CON-EVs or CCA-EVs were subjected to 12-15% SDS-PAGE for protein separation and Western blotting analysis. Coomassie blue gel staining was used as a loading control. Expression of sEV markers: CD81 (22 kDa), CD63 (28 kDa), Tsg101 (46 kDa), Flotillin-1 (48 kDa), HSP70 (70 kDa) and Alix (90kDa); lipoprotein marker: ApoA1 (25 kDa) and m/lEV markers: cytochrome c (12 kDa) and beta-actin (42 kDa) are shown in CCA-EVs *vs*. CON-EVs, with myotube (MT) lysates as a positive control. **(B)** Quantification of immunoblot analysis from panel A is represented here. Data were analyzed using an unpaired Student’s t-test, are expressed as scatter plots with mean (n=6-9). Exact p values for significant results (p<0.05) are shown.

**Figure 4.**
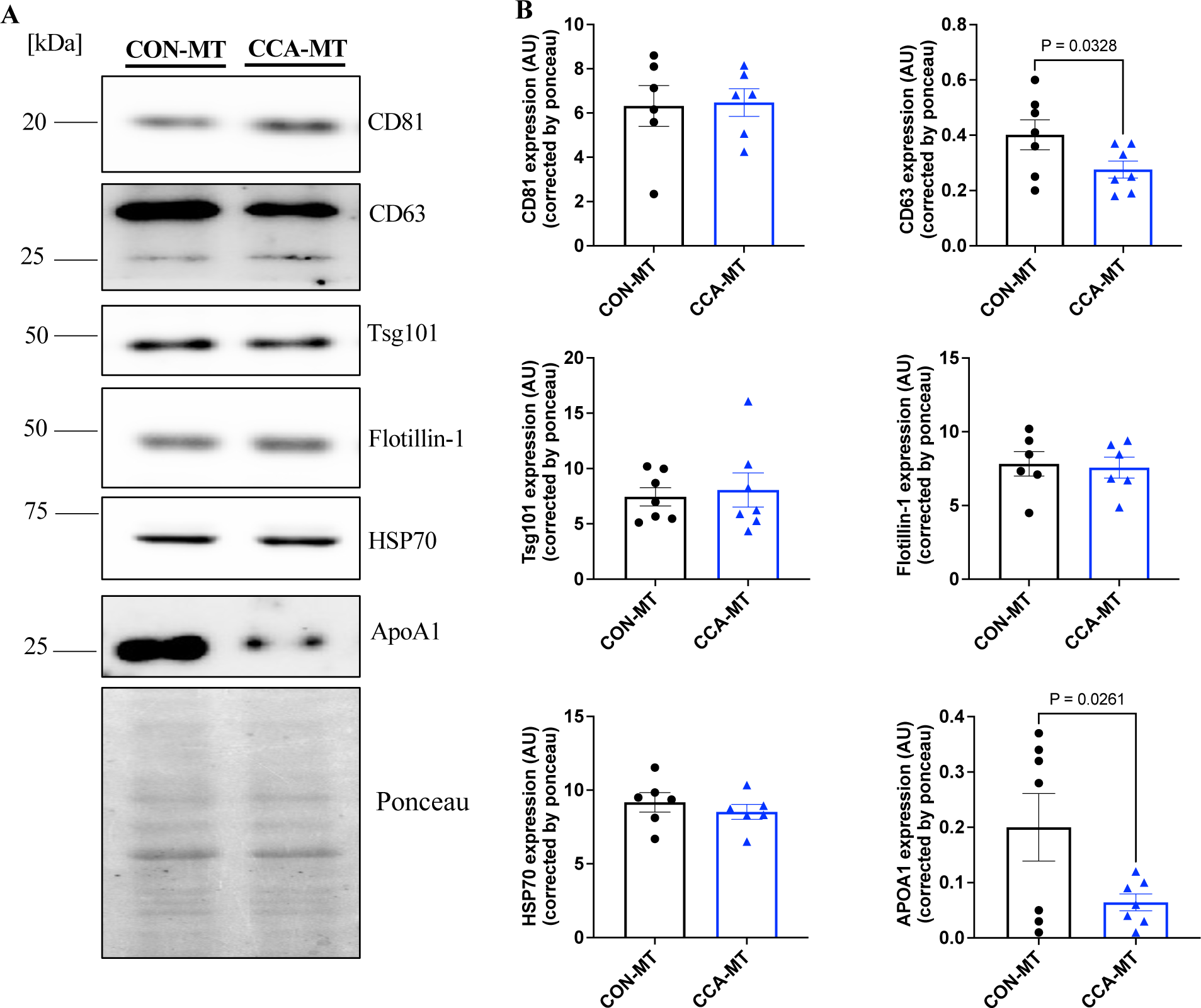
Effect of CCA on the expression of proteins related to EV biogenesis, cargo recruitment, and signalling in myotube (MT) lysates. **(A)** Equal amounts (5 µg) of proteins from CON-MT or CCA-MT extracts were resolved on a 12% SDS-PAGE for Western blotting analysis. β-Actin was used as a loading control. Protein expression of CD81, CD63, Tsg101, Flotillin-1, HSP70, and ApoA1 are shown in CCA-MT *vs*. CON-MT. **(B)** Quantification of immunoblot analysis from panel A is represented here. Data were analyzed using an unpaired Student’s t-test, are expressed as scatter plots with mean (n=6-7). Exact p values for significant results (p<0.05) are shown.

### CCA increases small EV secretion after each bout of contractile activity

We measured the effect of each bout of CCA on the biophysical properties of Skm-EVs as shown in the schematic (**Fig. 5A**) in tandem to determine temporal changes in EV properties with each day of contraction. While there was no difference in the average EV size between CCA *vs.* CON on any day, overall EV mean size on Day 4 was smaller compared to Day 1 (p=0.0349, n=4, **Fig. 5B**). EV concentration increased with CCA and with time (main effects using two-way ANOVA) with multiple comparison analysis showing that CCA-EVs had a 2.8-fold increase on Day 3 (p=0.0059, n=4) and a 1.5-fold increase on Day 4 (p=0.0151, n=4) compared to CON-EVs (**Fig. 5C**). The size/concentration distribution of EVs segregated by each day of CCA demonstrated an enrichment of sEVs in CCA-EVs *vs.* CON-EVs on each day (**Fig. 5D-G**). Zeta potential remained unchanged between groups and with time (**Fig. 5H**). Lastly, while total EV protein yield was not different between groups, it increased with time, with Day 4 EVs being higher than Day 1 EVs (p < 0.0001; n=4), *vs*. Day 2 EVs (p = 0.0019; n=4), and *vs*. Day 3 EVs (p=0.0032; n=4, **Fig. 5I**). Overall, our results demonstrate a temporal- and CCA-dependent increase in EV concentration, particularly sEVs, in tandem with progressively higher total EV protein yield with each day of CCA.

**Figure 5.**
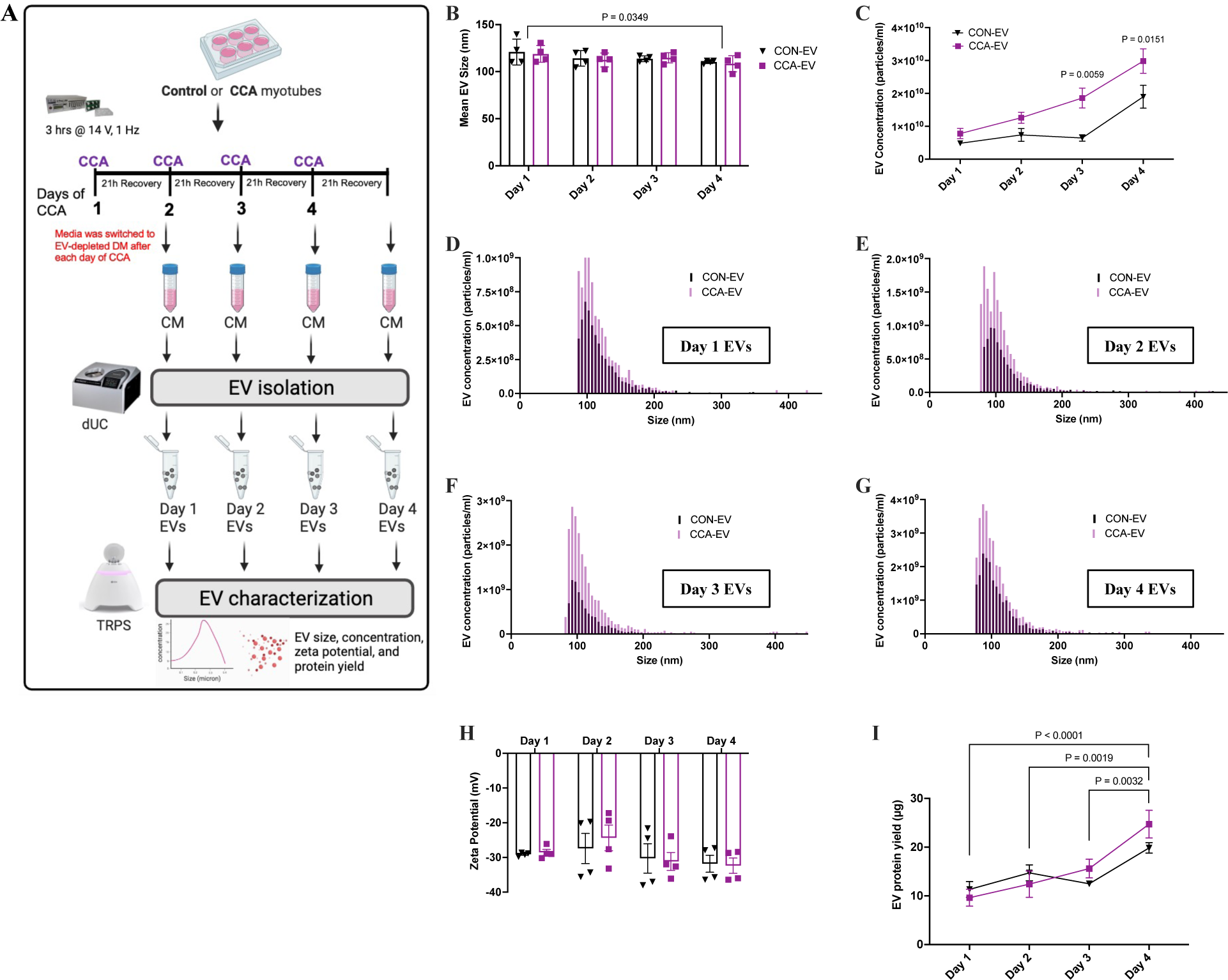
CCA enhances small EV secretion after each bout of contractile activity. (**A)** Myotubes were divided into CON and CCA plates. In the CCA group, myotubes were electrically paced for 3h/day at 14 V to mimic chronic endurance exercise *in vitro*. After the first bout of contractile activity, media was switched to exosome-depleted differentiation media in both CON and CCA groups, and myotubes were allowed to recover for 21 h. Conditioned media from control or stimulated myotubes was collected, and EVs were isolated via dUC. This process was done after each bout of contractile activity from Day 1-Day 4. EVs were characterized by size, zeta potential and concentration using TRPS. **(B)** Average EV size, **(C)** EV concentration, **(D-G)** EV size distribution segregated by each day of CCA (Day 1 – 4 EVs), **(H)** zeta potential and **(I)** EV protein yield in Day 1 -Day 4 EVs from CCA *vs*. CON groups. Panel C is showing significant differences between groups, i.e. CCA-EVs *vs*. CON-EVs, while panels B and I are demonstrating differences with time, i.e. between each day of contractile activity. Data were analyzed using a two-way ANOVA, with multiple comparisons corrected using Bonferroni’s post hoc test (n=4). Exact p values for significant (p<0.05) or trend to significant results are shown. Figure 5A created with BioRender.com.

### CCA-EVs do not affect cell count, viability, and protein yield in recipient skeletal muscle cells

Next, we investigated the functional effects of CCA-EVs on recipient myocyte growth including cell count, viability, and protein yield. 90,000 myoblasts/well was seeded in a standard 6-well plate and treated with Day 1 – 4 CON-EVs or CCA-EVs for 4 consecutive days as shown in the schematic (**Fig. 6A**). All EVs isolated each day from 10 ml of conditioned media were used to treat the cells, to recapitulate the physiological dosage of secreted EVs after each bout of contractile activity. There was no effect of EV treatment on cell count (**Fig. 6B**), or cell viability whether measured using trypan blue exclusion (**Fig. 6C**) or via the MTT assay (**Fig. 6D**). Protein yield of EV-treated myoblasts also remained unchanged (**Fig. 6E**). Removal of sEVs from the EV-depleted conditioned media was confirmed by measuring particle size distribution (**Fig. S3**), and the direct effect of conditioned media and EV-depleted media (negative control) on cell growth parameters assessed as well (**Fig. S4**). Treatment with EV-depleted media decreased cell count compared to (**Fig. S4A**), however, cell viability and protein yield remained unchanged (**Fig. S4B and S4C**). Overall, CCA-EVs did not affect cell count, viability, and protein yield in treated myocytes.

**Figure 6.**
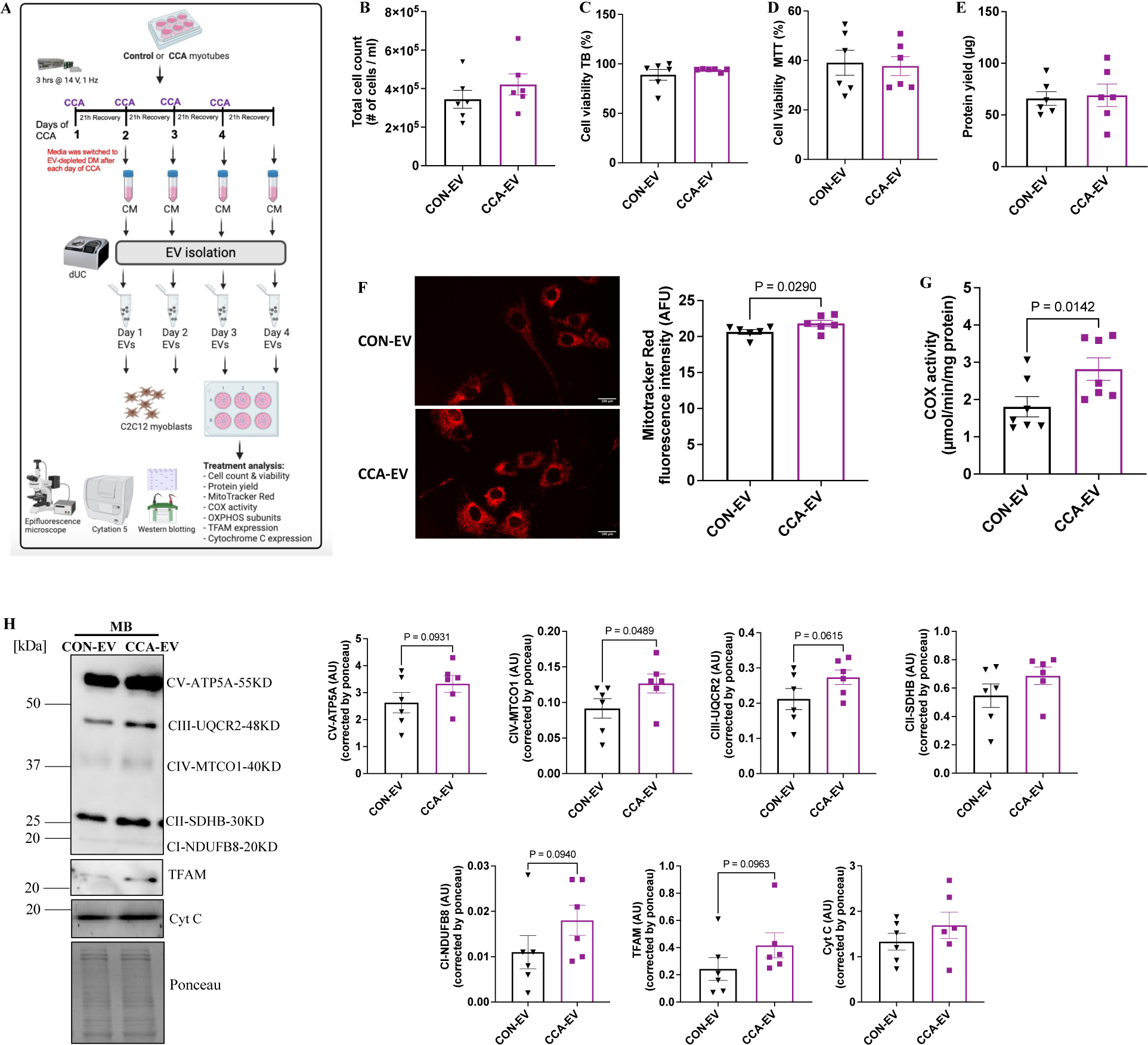
CCA-EV treatment did not affect cell count and viability, but increased mitochondrial biogenesis in recipient myoblasts (MB). **(A)** EVs were isolated after each bout of contractile activity as previously described, and co-cultured with myoblasts. After the last day of treatment, cells were collected and assessed for: **(B)** total cell count, **(C)** cell viability using trypan blue (TB), **(D)** cell viability using MTT, and **(E)** protein yield in myoblasts treated with CON-EVs or CCA-EVs. **(F)** Representative fluorescent images and quantification of MitoTracker Red staining in myoblasts treated with CON-EVs or CCA-EVs. Scale bar: 100 µm at 40X magnification. **(G)** COX activity, and **(H)** western blot analysis of OXPHOS subunits, TFAM and Cyt C in treated myoblasts. Data were analyzed using an unpaired Student’s t-test and expressed as scatter plots with mean (n=6). Exact p values for significant (p<0.05) or close to statistically significant results is shown. Figure 6A created with BioRender.com.

### CCA-EVs enhance mitochondrial biogenesis in recipient myocytes

To interrogate the effects of CCA-EVs on mitochondrial biogenesis, treated cells were stained with 50 nM of MitoTracker CMXRos and representative images at 40X showing mitochondrial content acquired for myoblasts treated with CON-EVs and CCA-EVs (**Fig. 6F**). We observed a 1.1-fold increase in MitoTracker red staining in myoblasts treated with CCA-EVs *vs.* CON-EVs (p=0.0290, n=6, **Fig. 6F**). This increase in MitoTracker Red staining was ameliorated when cells were treated with EV-depleted media treatment compared to conditioned media and EV treatment groups (**Fig. S4D**). Myoblasts treated with CCA-EVs displayed a 1.6-fold increase in COX activity than those treated with CON-EVs (p=0.0142, n=7 **Fig. 6G**). We measured the expression of select proteins associated with mitochondrial biogenesis, including OXPHOS protein subunits, TFAM and cytochrome *c*. We observed a 1.4-fold increase in the expression of electron transport chain subunit CIV-MTCO1 in myoblasts with CCA-EVs treatment (p=0.0489, n=6, **Fig. 6H**). The expression of other targets remained statistically unchanged with CCA-EV treatment, though some showed strong tendency towards an increase (**Fig. 6H**). Overall, our results indicate that prolonged CCA-EV treatment increased mitochondrial biogenesis in myoblasts.

### CCA-EVs enhanced mitochondrial oxygen consumption rates with pro-metabolic effect likely mediated by transmembrane and/or peripheral membrane proteins

Next, we treated myoblasts with CCA-EVs pretreated with proteinase K, Triton X-100 or the combination, and measured oxygen consumption rates (OCRs). Proteinase K will digest co-isolated proteins on the outside of EVs (EV corona) [48], as well as any transmembrane or peripheral membrane proteins decorating the outer surface of EVs since it cannot penetrate an intact membranous lipid bilayer. The addition of Triton X-100 will permeabilize the membrane, allowing all lumen and non-lumen EV proteins to be accessible for degradation by proteinase K. Thus, using proteinase K and Triton X-100 in combination with our EV treatment will allow us to determine if the pro-metabolic effects of CCA-EVs are likely mediated by EV corona, transmembrane and membrane-bound proteins *vs*. lumenal cargo proteins. A sample OCR tracing is shown for Seahorse XF Cell Mito Stress Test profile (Agilent Technologies) (**Fig. 7A**), and a representative OCR tracing is also shown for each condition in our study (**Fig. 7B**), with detailed quantification of basal respiration, maximal respiration, spare capacity, ATP production, protein leak and non-mitochondrial respiration rates. A 1.2-fold increase in maximal OCR with CCA-EV treatment *vs.* CON-EV was observed (p= 0.0171, n=8, **Fig. 7C**). Interestingly, this increase in maximal OCR was ameliorated by 1.4-fold when CCA-EVs were pretreated with proteinase K (p=0.0405 n=8, **Fig. 7C**), and remained reduced by a similar extent in Triton X-100 alone (p=0.0076, n=8, **Fig. 7C**), and in Triton X-100 + proteinase K combination groups (p=0.0171, n=8, **Fig. 7C**). A similar pattern was observed for spare capacity when CCA-EVs were subjected to proteinase K and Triton X-100 pre-treatment (**Fig. 7D**). ATP production increased with CCA-EV treatment compared to CON-EV treatment (p=0.0170, n=8, **Fig. 7E**). Basal respiration (**Fig. S5A**), proton leak (**Fig. S5B**) and non-mitochondrial respiration (**Fig. S5C**) rates were not affected by any treatment condition. Altogether, this indicates that CCA-EVs increase maximal OCR and ATP production in treated myoblasts, and this pro-metabolic effect is likely mediated by transmembrane and/or peripheral membrane proteins in CCA-EVs.

**Figure 7.**
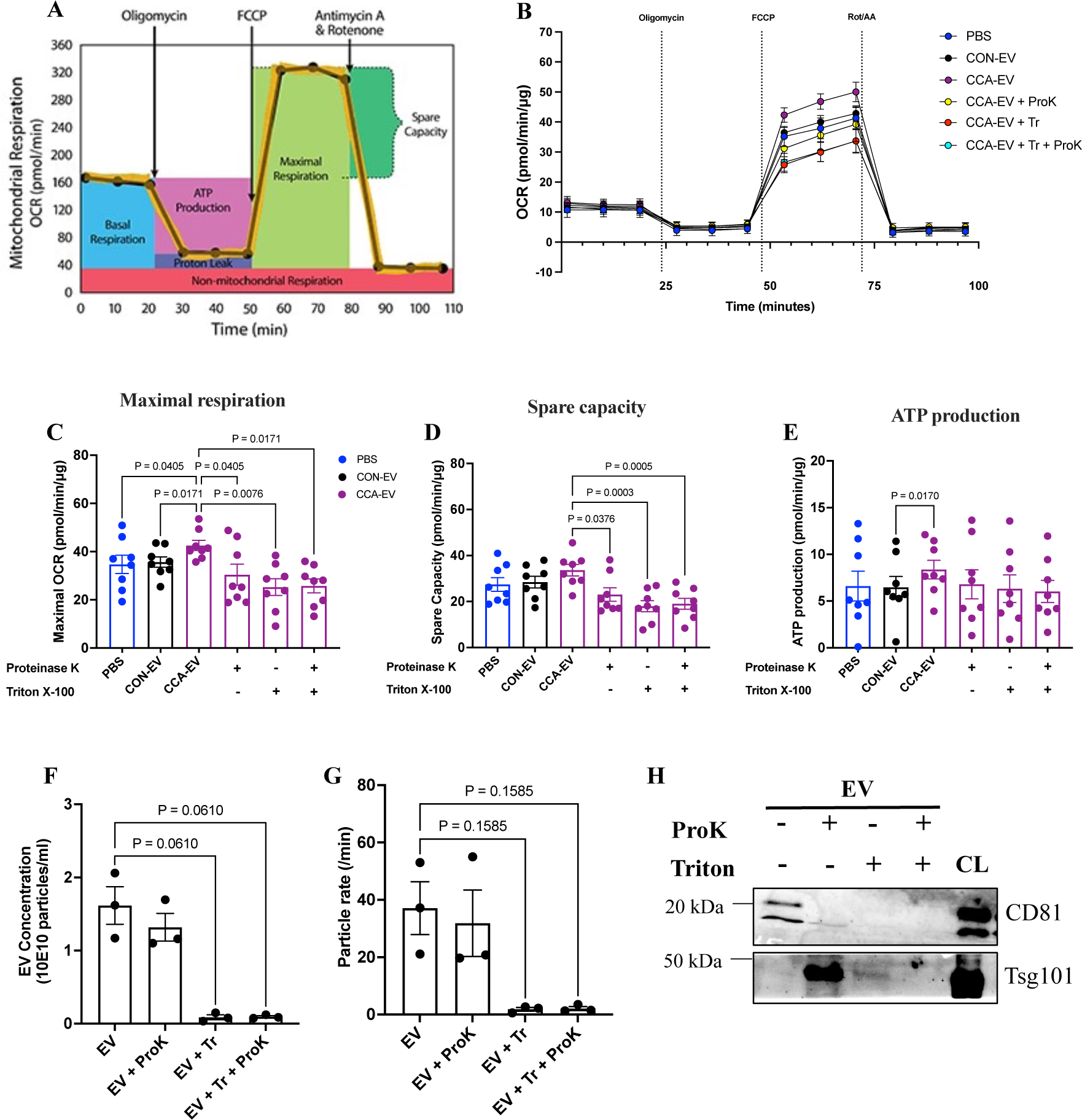
CCA-EVs increased oxygen consumption rates (OCR) in recipient myoblasts, likely mediated by EV transmembrane and/or peripheral membrane proteins. CCA-EVs were pretreated with or without 0.1% Triton X-100 and exposed to proteinase K (10 µg/mL, 1 h, 37 °C), and then co-cultured with MB for 4 days. MB were also treated with CON-EV and PBS for 4 days, after which OCR were measured. **(A)** Representative Seahorse XF Cell Mito Stress Test profile showing calculable parameters (Agilent Technologies). **(B)** Graphical representation of OCR for each group. **(C)** Maximal OCR, **(D)** spare (reserve) capacity, and **(E)** ATP production in treated myoblasts is shown. EVs were characterized by **(F)** concentration, **(G)** particle rate/min, and **(H)** expression of CD81 (transmembrane protein) and Tsg101 (lumenal cargo protein) in non-treated, vs. proteinase K ± Triton X-100 conditions. Data were analyzed using a one-way ANOVA, with multiple comparisons corrected using Holm-Šídák or Dunn’s post hoc test, and expressed as scatter plots with mean (n=3-8). Exact p values for significant (p<0.05) or close to statistically significant results are shown. CL – Cell lysates.

To ensure the effects of proteinase K on EV protein cargo in the presence or absence of Triton X-100 were as expected, we measured EV concentration and expression of transmembrane and lumenal cargo proteins. EV concentration reduced when treated with Triton X-100 with or without proteinase K (p=0.0610, n=3; **Fig. 7F**) compared with untreated EVs, indicative of EV membrane lysis. EV concentration in the proteinase K only treatment group remained unchanged when compared to untreated EVs (**Fig. 7F**). A similar pattern in particle rate was observed (**Fig. 7G**). Further, CD81 (EV transmembrane protein) expression was nearly abrogated when EVs were treated with proteinase K, Triton X-100 or the combination of both, but was present in the untreated group as well in the cell lysate fraction (**Fig. 7H**). Additionally, Tsg101 (EV lumen protein) content was unaffected after pre-treatment with proteinase K as expected, though decreased slightly in the Triton X-100 group, and was ameliorated in the combination treatment (**Fig. 7H**). These set of experiments confirmed the integrity of EV membranes in our EV preparation and that proteinase K only digested the proteins on the outside the EV membranes, unless used in combination with Triton X-100, as hypothesized.

### EV uptake by recipient myoblasts

Finally, to confirm uptake of EVs by recipient myocytes, we used the self-quenching membrane dye MemGlow 488 to label myotube-derived EVs and used two isolation methods, ultracentrifugation, and size exclusion chromatography (SEC), to remove free dye after which EVs were analyzed using flow cytometry. Using ultracentrifugation, we were able to retain 42.5% of MemGlow+ EVs (**Fig. 8A**), however SEC did not yield any MemGlow+ EVs (**Fig S6**). We performed the same steps using an equivalent volume of PBS with MemGlow dye, and unstained EVs as negative controls. There was no fluorescent signal in the unstained control, and only trace background fluorescent signal in the PBS control (**Fig. 8A**). Having established the specificity of MemGlow to label EVs, we next treated myoblasts with MemGlow+ EVs for 24 h, and evaluated MemGlow+ EV uptake in fixed EV-treated myoblasts using confocal microscopy (**Fig. 8B**). We observed the presence of green fluorescence indicative of MemGlow+ EVs, co-localizing with the red fluorescence from rhodamine phalloidin (F-actin) showing EV uptake in recipient myocytes (**Fig. 8B**). Similar results were obtained using PKH67 dye to monitor EV uptake (**Fig. 8C**).

**Figure 8.**
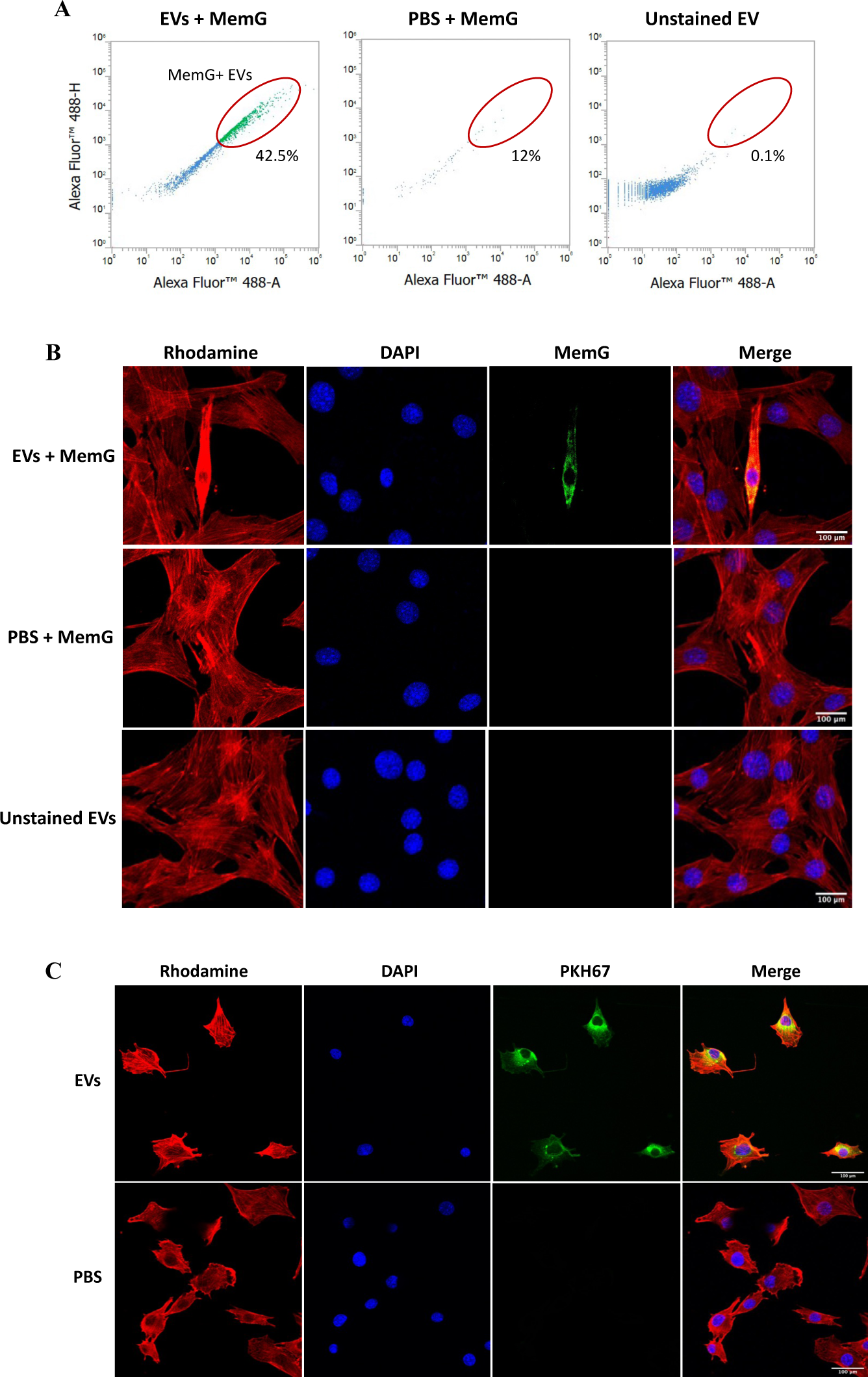
EV are taken up by recipient cells. **(A)** EVs isolated from conditioned media from myotubes or PBS were incubated with 200 nM MemGlow (MemG, green) for 1 h, then re-isolated with ultracentrifugation and measured by flow cytometry. Representative scatter plots are shown. **(B)** MemG-labelled EVs (green) or **(C)** 2 µM PKH67-labelled EVs (green) were used to treat myoblasts for 24 h, after which cells were fixed with 3.7% PFA, stained with Rhodamine phalloidin (red) and DAPI (blue), and imaged using confocal microscopy. PBS or unstained EVs were used as negative controls. Three representative images were taken per condition using the ZEN Pro software at 20X magnification (n=3), scale bar: 100 µm. Selected images demonstrate EV uptake by recipient cells.

## Discussion

Here we sought to elucidate the effect of chronic contractile activity (CCA) on the biophysical and functional properties of EVs released from skeletal muscle myotubes. To our knowledge, this is the first study to characterize EVs from skeletal muscle following chronic electrical pulse stimulation, a validated model of chronic exercise *in vitro* [36], [37]. The results demonstrate that chronic contractile activity of myotubes increased the secretion of EVs and in particular, the release of small EVs, enriched with sEV markers, Tsg101, CD81 and HSP70. CCA-EV treatment in myocytes increased mitochondrial biogenesis as determined by mitochondrial staining, enzyme assay, western blotting, and oxygen consumption analyses. Using proteinase K with or without Triton X-100, we determined that this pro-metabolic effect was likely mediated by transmembrane and/or peripheral membrane EV proteins. The specific protein(s) and underlying mechanism(s) that orchestrate these pro-metabolic adaptations have yet to be delineated.

Electrical pulse stimulation has been shown to effectively mimic the effects of chronic exercise on augmenting mitochondrial biogenesis *in vitro* [36], [49], an important adaptation to habitual endurance exercise. An increase in MitoTracker staining, COX activity and Cyt C (a critical protein in oxidative phosphorylation) in chronically stimulated myotubes, is in line with results reported by other groups [36], [37], [50], [51], and validated that the stimulation protocol as effective in mimicking the effects of CCA *in vitro*. We subsequently used it to interrogate the effects of contractile activity on Skm-EVs. TEM results showed that the purified EV preparations were enriched with vesicles between 30-150 nm in size, in line with previous work showing sEV release from C2C12 myotubes [52]–[55]. In this current study, we used TRPS, an orthogonal single-particle analysis technique to characterize polydisperse particles such as EVs from biological fluids [56], [57]. We found no difference in average EV size, but EV concentration increased in CCA-EVs *vs.* CON-EVs. EV size distribution results showed that CCA-EVs were enriched in sEVs compared to CON-EVs, and similar results were obtained using DLS. Zeta potential, an important indicator of colloidal stability of particles in dispersed systems [58], was not different between CCA-EVs *vs.* CON-EVs. These results indicate that chronic contractile activity of myotubes increase the release of sEVs. Interestingly, this is in line with other studies that have reported that endurance exercise training potentiates the release of sEVs in circulation [28], [31], [59].

Studies that have evaluated the effect of chronic exercise on EV concentration isolated EVs only after the whole exercise protocol was completed, and results can be confounded with just the “last bout effect” of exercise training intervention [28], [31], [59], [60]. Thus, to identify any temporal effects of each bout of contractile activity on the biophysical properties EVs, we isolated and characterized EVs after each day of CCA. We found that EV protein yield increased progressively with time, but not with contractile activity. On the other hand, the increase in EV concentration was both contractile activity- and time-dependent, in line with previous work that demonstrated 8 weeks of running in mice increased release of sEVs [31]. A significant increase in CCA-EV concentration *vs*. CON-EVs after Day 3 and Day 4 of contractile activity, could be as myotubes adapt to contractile activity and become “trained”, in agreement with metabolic capacity regulating EV secretion [35]. It will be interesting to measure the release of EVs in trained *vs*. untrained muscle in human or murine models *in vivo*. Altogether, our results showed that chronic contractile activity enhanced EV release, and this occurs mostly on Day 3 and Day 4 of contractile activity. While the progressive increase in total protein content of EVs is likely due to the concomitant elevation in sEV secretion from contracted myotubes, there might be protein-specific modifications in the EV cargo instead of just bulk elevation in total protein, though this needs to be established.

To ascertain the subtype of EVs isolated, we measured the expression of proteins including exosomal/sEV markers (CD81, CD63, Tsg101, HSP70, Alix and Flotillin-1), m/l EV markers (Cyt C and β-actin), and a non-EV marker (Apo-A1) as per MISEV guidelines [3]. Our results demonstrate that CCA-EVs are enriched with CD81, Tsg101 and HSP70 compared to CON-EVs, in line with previous studies that have shown that both acute and chronic exercise training increases EV protein levels of Tsg101, CD81 and HSP70 in mice and humans [24], [27], [31], [61], [62]. We did not detect any expression of Cyt C and β-actin in the EV lysates, likely as our EV preparations are enriched with sEVs. Cyt C (a mitochondrial protein) has been extensively used as a m/l EV marker [63], [64], and β-actin (a protein associated with the cytoskeleton) has been shown to be enriched in larger sized vesicles [65], [66]. To assess the purity of our Skm-EV isolation protocol, we measured the levels of ApoA1, a non-EV lipoprotein marker. It is not surprising that our EV preparation contains ApoA1 since we observed its expression in our myotube lysates, in agreement with previous reports [67]. Additionally, lipoproteins can be co-isolated with sEVs due to the overlap in their size and density [68], [69]. To reduce the level of lipoprotein contamination, it is advised that researchers use two or more isolation or purification steps, which is an approach we can utilize in future. Notwithstanding, both CON-EVs and CCA-EVs expressed ApoA1, and the expression of this non-EV co-isolate protein was not different between groups. Next, we determined if chronic contractile activity altered the expression of EV-related proteins in myotubes. We found that CCA myotubes expressed reduced ApoA1 and CD63 protein content compared to CON myotubes. However, we did not find a difference in CD81, Tsg101, HSP70 and Flotillin-1 between the groups, which was counter-intuitive since we observed an increase in CD81, Tsg101, and HSP70 in EVs from CCA myotubes. Nevertheless, our results agree with a recent study that showed no difference in CD81 and Tsg101 protein expression in skeletal muscle from healthy humans with acute exercise, even though these proteins were increased in circulatory EVs with exercise [62]. Altogether our results support the conclusion that Skm-EVs are enriched with sEVs, and could be of ectosomal or endosomal origin. Future experiments to investigate the mechanism(s) of EV biogenesis and secretion from skeletal muscle cells after chronic contractile activity are warranted.

Chronic exercise-derived EVs have been shown to delay the progression of prostate cancer [25], and induce an exercise intensity-dependent protective effect against hypoxia-induced apoptosis in endothelial cells [59]. EVs in circulation largely originate from endothelial cells, platelets, erythrocytes, leukocytes, [24], [27], as well as skeletal muscle, albeit at very low levels (∼5% of total EVs) [21]. While, it has been hypothesized that Skm-EVs may mediate the pro-metabolic effects of exercise [20], [70], the hypothesis has been largely untested due to the aforementioned challenges in purifying Skm-EVs from contracting muscle tissue. Here we present the first specific functional assessment of Skm-EVs obtained from contracting myotubes, without the confounding presence of non-Skm-EVs. To evaluate the biological functional activity of Skm-EVs isolated post-chronic contractile activity on recipient myocytes, we treated myoblasts with CON-EVs or CCA-EVs isolated after each bout of contractile activity for 4 days. Using MemGlow 488 and PKH67, we confirmed that EVs can be taken up by recipient myoblasts. CCA-EV co-culture treatment did not alter recipient cell count, viability, and protein yield compared to those treated with CON-EVs. We have previously shown that extracellular particles (EP) isolated following acute contractile activity increased cell count but not viability in myoblasts, but that study was done with a different exercise regimen, isolation method, and treatment protocol [42]. Another study reported that chronic exercise-derived EVs increased cell viability in Neuro2A cells, a mouse neuroblastoma cell line, also using different exercise regimen and protocols [71]. Interestingly, EV-depleted media treatment reduced cell count compared to conditioned media and CCA-EV treated cells, illustrating the EV-dependent nature of these adaptations, and in concordance with previous work [42]. Similarly, EV depletion mostly from serum (fetal bovine serum or horse serum) has been reported to stunt cell growth [72], [73]. This indicates that EVs are important for cell growth and viability.

Increased mitochondrial content and function represents a key adaptive response to sustained contractile activity/exercise. The initiation of exercise-induced mitochondrial biogenesis begins with the onset of contractile activity which induces a series of molecular events leading to the upregulation of PGC-1α, and a subsequent activation of nuclear respiratory factor 1 (NRF-1) and mitochondrial transcription factor A (TFAM) [74]. This promotes a coordinated downstream increase in both nuclear and mitochondrial proteins involved in mitochondrial biogenesis [74]. Our results showed that mitochondrial biogenesis as measured by MitoTracker staining, COX activity and expression of proteins related to mitochondria (e.g., complex IV subunit I CIV-MTCO1) was increased in myoblasts treated with CCA-EVs. We observed a 1.7-fold increase in TFAM expression with CCA-EV treatment, which although not statistically significant, in tandem with the increase in complex IV subunit I could indicate an induction of the de novo mitochondrial biogenesis pathway. Additionally, mitochondrial biogenesis can be induced by modifications in mitochondrial dynamics (movement, speed) and regulation of mitochondrial quality control (fusion and fission) [75]. We do not know if CCA-EV treatment activated these alternate pathway(s). It is important to note that we treated recipient cells with the total EVs isolated after each day of contractile activity to recapitulate the physiological relevance of secreted EVs isolated after each bout of exercise. This poses the question whether the significant effects observed with CCA-EV treatment on myoblasts are due to the increased EV concentration from stimulated myotubes, or because of changes in EV cargo, or both. This warrants future work to evaluate the dose effect of CCA-EVs.

To evaluate the specificity of the pro-metabolic effect of CCA-EVs further, we co-cultured myocytes with CCA-EVs pre-treated with proteinase K, Triton X-100 or both, and measured oxygen consumption rates. While proteinase K digests proteins, Triton X-100 permeabilizes EV membranes [76], resulting in the corollary loss of membrane proteins. In concordance with the putative effects of proteinase K and Triton X-100, Triton X-100 decreased EV concentration and particle rate. The proteinase K only treated group had similar concentration and particle rate as the untreated group, as proteinase K digestion is limited to the membrane and extra-membranous (EV corona) proteins if the purified EVs are structurally intact. CD81, a membrane tetraspanin, disappeared with proteinase K treatment alone, as well as with Triton X-100 only, and in the combination group. Previous studies showed similar result for CD81and CD63 after EVs were treated with proteinase K alone [77], [78], [79], but that CD63 expression remains unaltered with Triton X-100 only [79]. The decrease in CD81 after Triton X-100 treatment in our study could be due to the loss of fractured EV membranes (with embedded proteins) that pelleted during the additional ultracentrifugation step introduced post-treatment with proteinase K and Triton X-100. This extra ultracentrifugation step was necessary as Triton X-100 would kill recipient cells if not properly neutralized. The expression of Tsg101, an EV lumen cargo protein, was ameliorated after co-treatment of EVs with Triton X-100 and proteinase K, in agreement with previous work [79].

CCA-EVs significantly elevated maximal respiration rates, and ATP production *vs.* CON-EV in myocytes. Notably, the CCA-EV induced increase in maximal respiration was completely abrogated with proteinase K pre-treatment of EVs, illustrating the importance of transmembrane, peripheral membrane-bound and/or EV corona proteins in perpetuating CCA-evoked pro-metabolic effects. The maximal respiration rates remained reduced in the Triton X-100-treated CCA-EVs, likely due to loss of membrane proteins due to membrane disruption. The combination of Triton X-100 and proteinase K did not cause an additive decrement in respiration *vs*. Triton X-100 only group. This could indicate the EV lumen proteins are not as important in mediating the metabolic adaptations. These findings concur with recent reports regarding removal of surface proteins from EVs by proteinase K and the subsequent attenuation in EV functional effects [77]. Additionally, it is likely that transmembrane and peripheral membrane proteins in CCA-EVs are necessary for EV internalization through multiple pathways, and/or critical for cell surface receptor-mediated signaling downstream. The removal of these membrane proteins may thus block activation of signaling pathways, and/or delivery of EV cargo to recipient cells. Further research to elucidate the mechanism(s) involved is clearly needed.

In summary, the CCA model we used in this study helps to circumvent the challenge of using whole blood-based EV preparations to study the functional role of Skm-EVs. CCA is an efficient, fast method to induce mitochondrial biogenesis in cells, in as quickly as four days, and to interrogate the effect of prolonged contractile activity on Skm-EVs secretion, properties and downstream biological activity, bypassing the limitations of using circulatory EVs. Our findings illustrate that CCA induced the preferential release of sEVs, significantly enriched with Tsg101, CD81 and HSP70. We also showed for the first time that Skm-EVs isolated post-CCA increased mitochondrial biogenesis in myoblasts increasing both mitochondrial content and function, and this effect is likely mediated by the transmembrane and peripheral membrane-bound EV proteins. The specific EV protein(s) and underlying mechanism(s) involved have yet to be elucidated. These novel hitherto unreported findings in this manuscript demonstrate the ability of CCA-EVs to transmit pro-metabolic signaling, recapitulating the effect of contractile activity in healthy myocytes. Whether a similar effect is observed in other cell types, and *in vivo*, remains to be ascertained.

## Author Contributions

P.O.O., T.S.G., S.S., N.K. and B.B. performed experiments in the current study; P.O.O analyzed data, created figures, and prepared manuscript. A.R.W. and J.W.G. provided technical and theoretical expertise to complete the work. A.S. designed the project, and helped analyze data, create figures, write, and edit the manuscript. A.S. is the corresponding author and directly supervised the project. All authors have read, edited and agreed to the published version of the manuscript.

## Acknowledgements

The authors would like to thank Dr Hagar Labouta, College of Pharmacy, Rady Faculty of Health Sciences at the University of Manitoba for assistance with the EV characterization experiments using dynamic light scattering (DLS); Martha Hinton for help with the EV uptake experiments using confocal microscopy; Emily Turner-Brannen for providing assistance with the COX activity experiment using Agilent BioTek Cytation; and Taiana M. Pierdoná for the technical support during this project.

## Conflict of Interest

All other authors declare no conflict of interest. The funders had no role in the design of the study; in the collection, analyses, or interpretation of data; or in the writing of the manuscript.

Word count (not including title page, abstract, methods, references, figure legends):

