## supplementary files for "Chronic contractile activity induced skeletal muscle-derived extracellular vesicles increase mitochondrial biogenesis in recipient myocytes via transmembrane or peripheral membrane proteins"

### Supplementary Methods

#### Dynamic light scattering (DLS)

Measurements of EV size and zeta potential were performed by phase analysis DLS using a NanoBrook ZetaPALS (Brookhaven Instruments, Holtsville, NY, USA) in collaboration with Dr. Hagar Labouta's lab (College of Pharmacy, University of Manitoba). EV samples were diluted 1:75 with 0.22 μm filtered PBS. Five measurements were recorded for each sample with a dust cut-off set to 40. EV size was measured as an intensity averaged multimodal distribution using a scattering angle of 90°, and size bins were used to represent total size intensity within a given size range. Zeta potential analysis was performed using a Solvent-Resistant Electrode (NanoBrook) and BI-SCGO 4.5 mL cuvettes (NanoBrook). Smoluchowski formula was used to calculate zeta potential and final results were averaged irrespective of the charge (negative/positive) [80]. All measurements were performed in PBS (pH 7.4) at 25 °C.

A

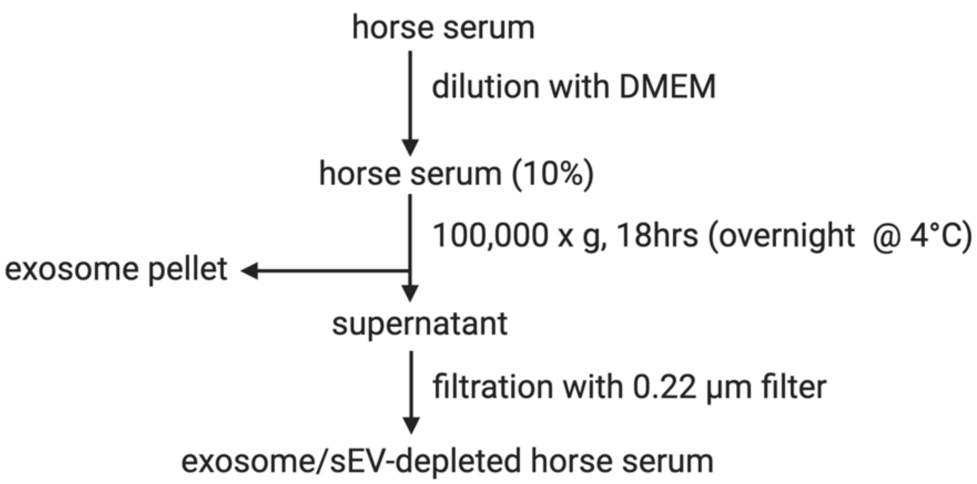


B

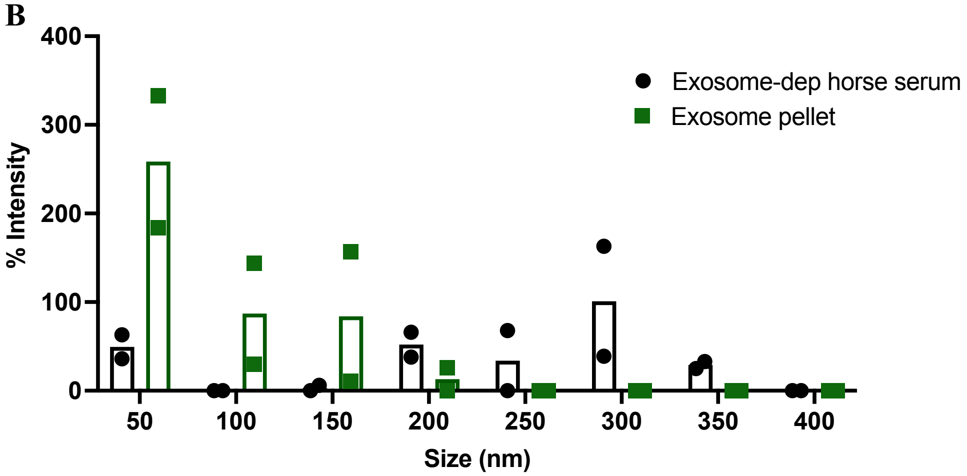


**Figure S1. Characterization of exosome-depleted horse serum.** (**A**) Flow chart demonstrating the horse serum exosome-depletion protocol. **(B)** EV size distribution (measured using DLS) illustrating nanovesicles captured in exosome-depleted horse serum when compared to the exosome pellet after the 100,000*x*g, 18hr spin (n=2).

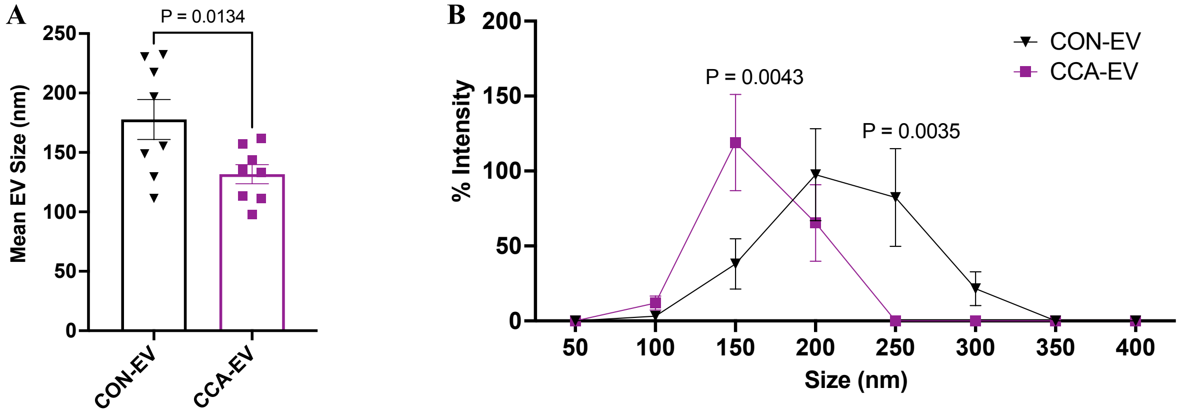


**Figure S2. Effect of CCA on the biophysical properties of EVs using DLS. (A)** CCA was done as previously described. Conditioned media from control or stimulated myotubes was collected. EVs were isolated via differential ultracentrifugation (dUC) and characterized by size and concentration using dynamic light scattering (DLS; NanoBrook ZetaPALS). **(A)** Average EV size, and **(B)** EV size distribution in CCA-EVs *vs*. CON-EVs. Data were analyzed using an unpaired Student's t-test in panel A, and by a two-way ANOVA in panel B, with multiple comparisons corrected using Bonferroni's post hoc test (n=8). Exact p values for significant results ($p<0.05$) are shown.

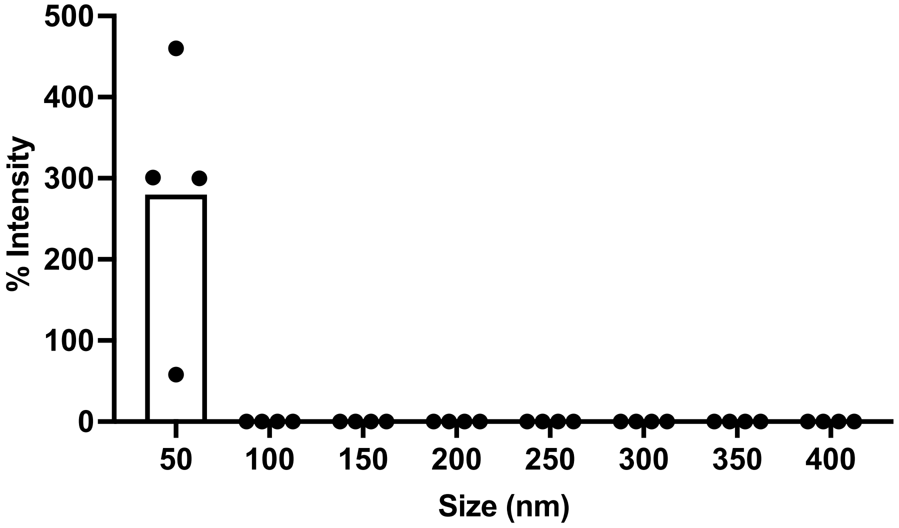


**Figure S3. Characterization of EV-depleted conditioned media.** Size distribution analysis of nanoparticles (measured using DLS) in the EV-depleted conditioned media after the first 100,000*x*g spin during dUC (n=4).

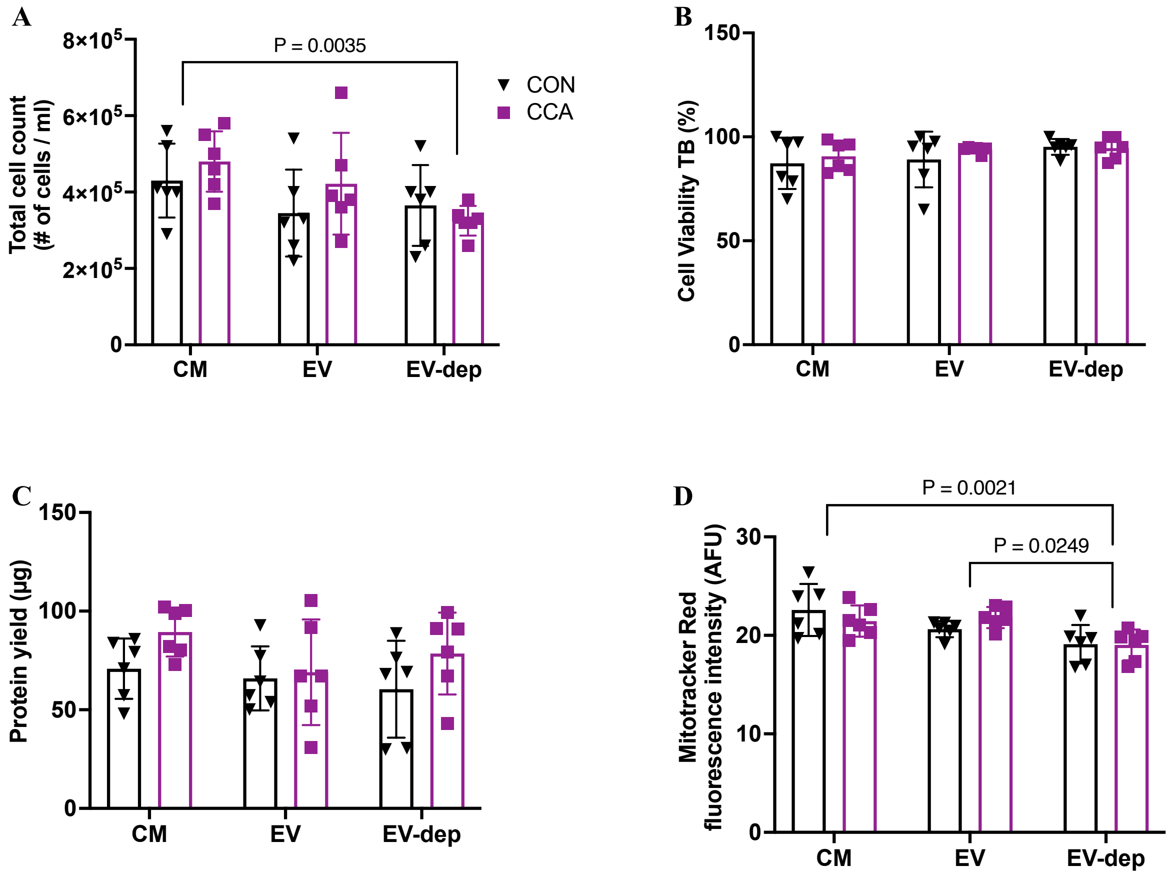


**Figure S4. Effect of CM, EV and EV-dep treatment on myoblasts.** MB were co-cultured with either conditioned media (CM), EVs or EV-depleted conditioned media (EV-dep) from CON-MT or CCA-MT for 4 days after each bout of contractile activity. After treatment, markers for cell count, viability and mitochondrial content were measured. (**A)** Total cell count, **(B)** Cell viability with trypan blue (TB), **(C)** Protein yield and **(D)** MitoTracker Red staining in treated myoblasts with CM, EV and EV-dep treatment. Graphs A and D are showing significant differences between treatment groups. Data were analyzed using a two-way ANOVA, with multiple comparisons corrected using Bonferroni's post hoc test (n=6). Exact p values for significant ($p<0.05$) results are shown.

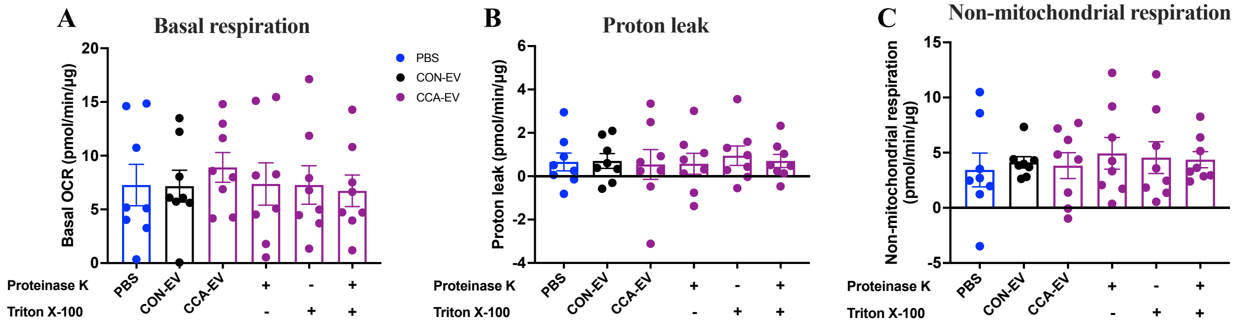


**Figure S5. Effect of CCA-EV treatment on basal and non-mitochondrial respiration, and proton leak.** CCA-EVs were pretreated with or without 0.1% Triton X-100 and exposed to proteinase K (10 µg/mL, 1 h, 37 °C), and then co-cultured with MB for 4 days. MB were also treated with CON-EV and PBS for 4 days, after which mitochondrial respiration was measured. **(A)** ATP production**, (B)** Proton leak, and **(H)** non-mitochondrial respiration in treated myoblasts is shown. Data were analyzed using a one-way ANOVA, with multiple comparisons corrected using Holm-Šídák or Dunn's post hoc test, and expressed as scatter plots with mean (n=8). No statistically significant results were observed with any treatment condition.

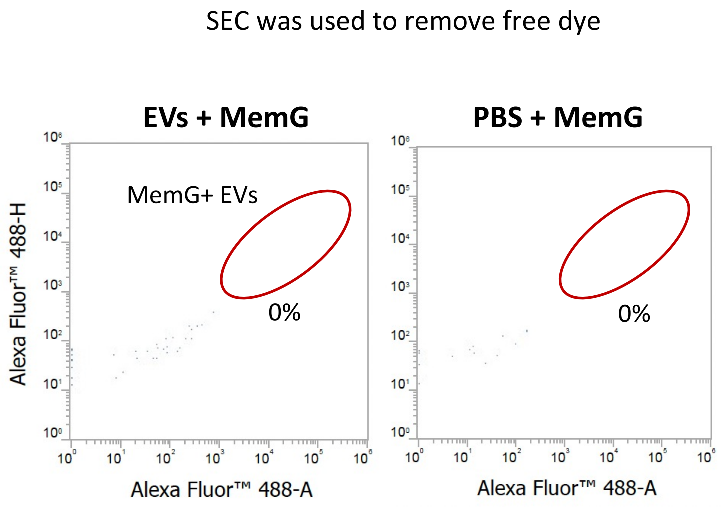


**Figure S6. Using SEC to remove free MemGlow 488**. EVs or PBS were incubated with MemG (green) for 1 h, then re-isolated with SEC and measured by flow cytometry.
